# Microbial metabolite butyrate-prodrug polymeric micelles demonstrate therapeutic efficacy in pre-clinical models of food allergy and colitis

**DOI:** 10.1101/2022.05.01.490224

**Authors:** Ruyi Wang, Shijie Cao, Mohamed Elfatih H. Bashir, Lauren A. Hesser, Yanlin Su, Sung Min Choi Hong, Andrew Thompson, Elliot Culleen, Matthew Sabados, Nicholas P. Dylla, Evelyn Campbell, Riyue Bao, Eric B. Nonnecke, Charles L. Bevins, D. Scott Wilson, Jeffrey A. Hubbell, Cathryn R. Nagler

## Abstract

The microbiome modulates host immunity and aids in maintenance of tolerance in the gut, where microbial and food-derived antigens are abundant. Modern lifestyle practices, including diet and antibiotic use, have depleted beneficial taxa, specifically butyrate-producing Clostridia. This depletion is associated with the rising incidence of food allergy, inflammatory bowel diseases, and other noncommunicable chronic diseases. Although butyrate is known to play important roles in regulating gut immunity and maintaining epithelial barrier function, its clinical translation is challenging due to its offensive odor and quick absorption in the upper gut. Here, we have developed two polymeric micelle systems, one with a neutral charge (NtL-ButM) and one with a negative charge (Neg-ButM) that release butyrate from their polymeric core in different regions of the gastrointestinal tract when administered intragastrically to mice. We show that these butyrate-containing micelles, used in combination, restore a barrier-protective response in mice treated with either dextran sodium sulfate or antibiotics. Moreover, butyrate micelle treatment protects peanut-allergic dysbiotic mice from an anaphylactic reaction to peanut challenge and rescues their antibiotic-induced dysbiosis by increasing the abundance of *Clostridium* Cluster XIVa. Butyrate micelle treatment also reduces the severity of colitis in a murine model. By restoring microbial and mucosal homeostasis, these butyrate-prodrug polymeric micelles may function as a new, antigen-agnostic approach for the treatment of allergic and inflammatory disease.

## Introduction

The gut microbiome has many effects on both mucosal and systemic health^1–3^. While many mechanisms by which bacteria regulate mucosal homeostasis remain unknown, short-chain fatty acids, especially butyrate, have been well studied as immunoregulatory molecules^4–5^. Butyrate is produced by a subset of intestinal bacteria through the fermentation of dietary fiber^6^. Decreased abundance of butyrate-producing bacteria has been observed in human cohorts of food allergy, asthma, inflammatory bowel disease (IBD), and other noncommunicable chronic diseases (NCCDs)^7–13^. However, oral delivery of butyrate to the small intestine (where food antigens are absorbed) and large intestine (where most commensal bacteria reside) has been a challenge.

Butyrate, even with enteric coating or encapsulation, possesses a foul and lasting odor and taste. As a sodium salt, orally administered butyrate is not absorbed in the part of the gut where it can have a therapeutic effect and is metabolized too rapidly to maintain a pharmacologic effect^14^. Previous work in murine models that demonstrated therapeutic effects of butyrate relied on high concentration, *ad libitum* exposure to butyrate (mM quantities in drinking water for several weeks) or utilized butyrylated starches^15–21^. In human trials, intrarectal delivery of butyrate is moderately efficacious in treating colitis but is not a preferred route of administration^22^. A more controlled and practical delivery strategy is needed to exploit the potential therapeutic benefits of butyrate clinically to treat allergic and inflammatory diseases of the lower gastrointestinal (GI) tract. To address this problem, we designed block copolymers that can form water-suspensible micelles carrying a high content of butyrate in their core. These polymer formulations mask the smell and taste of butyrate and act as carriers to release the active ingredient (butyrate) over time as the micelles transit the GI tract. They can be formulated and administered as a suspension, allowing high dose without requiring a large number of pills. We developed two novel polymers: one with a neutral charge which predominantly releases butyrate in the ileum (NtL-ButM), and one with a negative charge which predominantly releases butyrate in the cecum (Neg-ButM). This micelle system allows us to deliver butyrate to its site of biological effect and overcome the existing limitations of oral administration.

Butyrate is produced by certain members of the Clostridia class in the distal GI tract and is the preferred energy substrate for colonic epithelial cells, strengthens gut barrier function by stabilizing hypoxia-inducible factor and maintains epithelial tight junctions^5,23^. To mediate its immunomodulatory functions, butyrate acts via signaling through specific G protein coupled receptors or as an inhibitor of histone deacetylase activity (HDACs)^24^. HDAC inhibition by SCFAs promotes the differentiation of colonic regulatory T cells (Tregs)^17–19^. The ability of butyrate to regulate both barrier and adaptive immunity makes it an ideal drug candidate for inflammatory diseases such as food allergy and colitis, which are mediated by both impaired intestinal barrier function and effector T cells.

We have previously shown a protective role for butyrate-producing Clostridia in prevention of food allergies in both mouse models and human cohorts^7,8,10,25^. Neonatal administration of antibiotics reduces intestinal microbial diversity and impairs epithelial barrier function, resulting in increased access of food allergens to the systemic circulation^25^. Administration of a consortium of spore-forming bacteria in the Clostridia class restored the integrity of the epithelial barrier and prevented allergic sensitization to food^25^. We went on to demonstrate a causal role for bacteria present in the healthy infant microbiota in protection against cow’s milk allergy^8^. Transfer of the microbiota from healthy, but not cow’s milk allergic (CMA), human infants into germ-free (GF) mice protected against an anaphylactic response to a cow’s milk allergen. By integrating differences in the microbiome signatures present in the healthy and CMA microbiotas with the changes each induced in ileal gene expression upon colonization of GF mice, we identified a single butyrate producing Clostridial species, *Anaerostipes caccae*, that mimicked the effects of the healthy microbiota upon monocolonization of GF mice^8^. Recent findings from a diverse cohort of twin children and adults concordant and discordant for food allergy validated the mouse model data with human microbiome samples. We found that most of the operational taxonomic units (OTUs) differentially abundant between healthy and allergic twins were in the Clostridia class; the broad age range of the twins studied indicated that an early-life depletion of allergy-protective Clostridia is maintained throughout life^10^.

Like food allergy, the incidence of IBD, including ulcerative colitis and Crohn’s Disease, has been rapidly increasing and is dependent on the commensal microbiota^11,26^. Bacterial dysbiosis in patients with IBD has been well characterized and consistently shows decreased abundance of butyrate-producing species^12,13^. Clinical trials have attributed the efficacy of fecal transplant for the treatment of *Clostridium difficile* infection and ulcerative colitis to some engraftment of butyrate-producing taxa including *Lachnospiraceae and Ruminococcaceae (Clostridium* clusters XIVa and IV)^27,28^. However, long-term engraftment of oxygen-sensitive anaerobic bacteria from FMT or oral delivery of consortia has proven challenging^29,30^. We therefore sought to explore butyrate itself as a candidate drug to maintain both microbial and mucosal homeostasis in the dysbiotic gut and to treat the consequences of inappropriate responses to the contents of the gastrointestinal lumen, including both dietary antigens (food allergies) and bacterial products (IBD).

The butyrate-conjugated polymer formulations developed herein release butyrate in distinct segments of the lower GI tract, in contrast to sodium butyrate (NaBut), which is predominantly absorbed in the stomach. When the two polymers are administered together (ButM), they modulate barrier integrity in antibiotic-treated mice and in mice treated with dextran sodium sulfate (DSS), a chemical perturbant that induces epithelial barrier dysfunction. Intragastric administration of our butyrate-prodrug micelles ameliorates an anaphylactic response to peanut challenge in a mouse model of peanut allergy with dysbiosis. To our knowledge this is the first evidence that butyrate can effectively treat previously established food allergy in a pre-clinical model. We hypothesize that this therapeutic effect may be elicited in part by modulating the microbiome, as treatment with ButM increases the abundance of bacteria in Clostridial clusters IV and XIVa known to contain butyrate-producing taxa. Moreover, treatment with ButM reduces disease severity in a T cell transfer model of colitis. These findings pave the way for future clinical translation of butyrate micelles in treating food allergies and inflammatory bowel disease.

## Results

### Co-polymers formulate butyrate into water-suspensible micelles

The block copolymer amphiphile pHPMA-b-pBMA was synthesized through two steps of reversible addition-fragmentation chain-transfer (RAFT) polymerization (**Fig. 1a**). The hydrophilic block was formed from N-(2-hydroxypropyl) methacrylamide (HPMA), while the hydrophobic block was from N-(2-butanoyloxyethyl) methacrylamide (BMA), thus connecting a backbone sidechain to butyrate with an ester bond. This ester bond can be hydrolyzed in the presence of digestive esterases and releases butyrate in the GI tract, resulting in a water-soluble polymer as a final product. In addition to pHPMA-b-pBMA, we also synthesized pMAA-b-pBMA, which has an anionic hydrophilic block formed from methacrylic acid (MAA) (**Fig. 1a**). At the block size ratios used herein, both pHPMA-b-pBMA and pMAA-b-pBMA contain 28% of butyrate by weight.

**Fig. 1.**
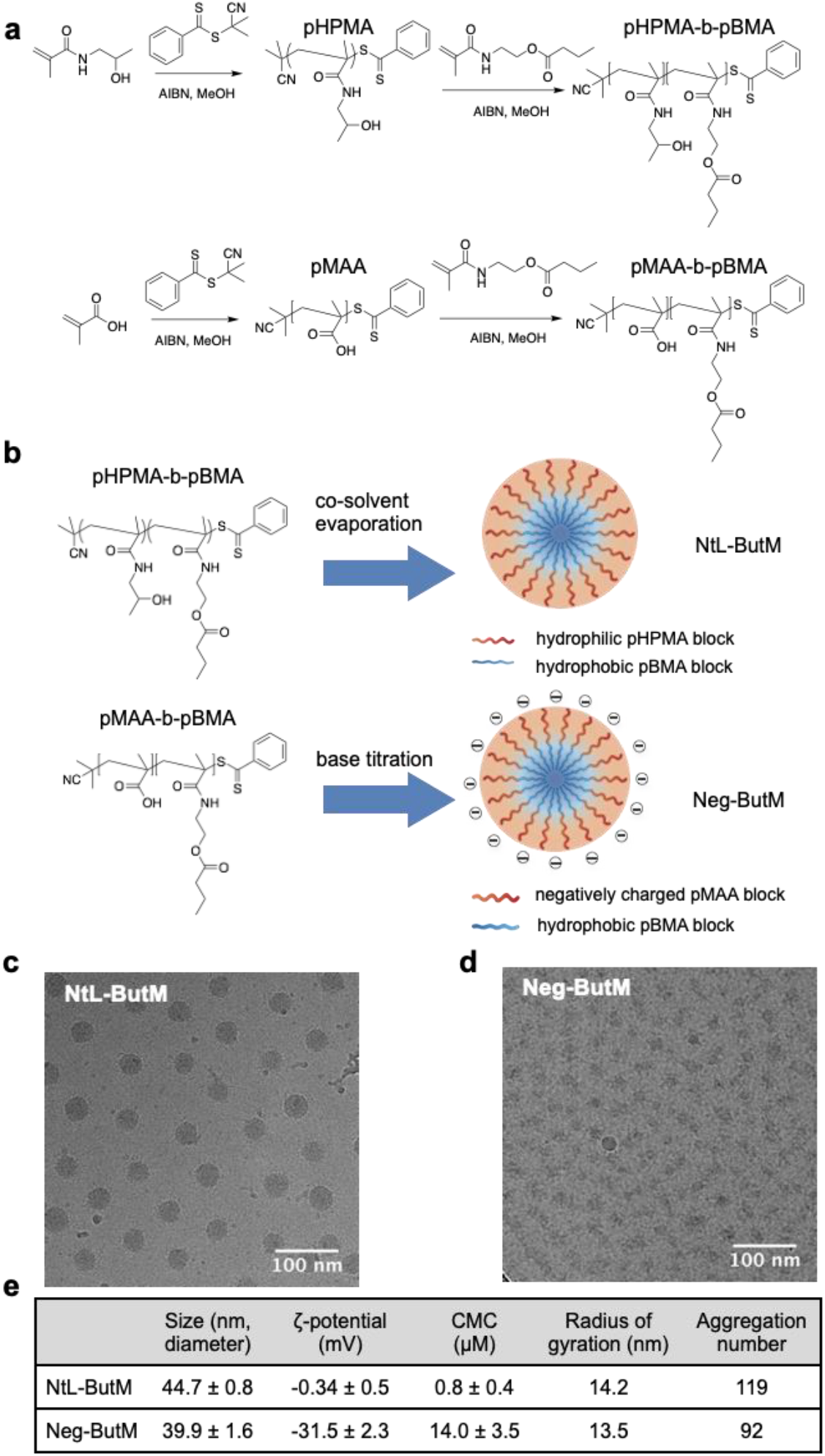
Chemical composition and structural characterization of butyrate-prodrug micelles, namely NtL-ButM, consisting of the neutral block copolymer pHPMA-b-pBMA, and Neg-ButM, consisting of the anionic block copolymer pMAA-b-pBMA. **a,** Synthetic route of pHPMA-b-pBMA and pMAA-b-pBMA. **b,** (**upper**) The NtL-ButM contains a hydrophilic (HPMA) block as the micelle corona, while a hydrophobic (BMA) block forms the micelle core. (lower) The Neg-ButM contains a hydrophilic (MAA) block that forms a negatively charged micelle corona, and the same hydrophobic (BMA) block as NtL-ButM. **c, d,** Cryogenic electron microscopy (CryoEM) images show the spherical structures of micelles NtL-ButM (**c**) or Neg-ButM (**d**). **e,** Table summarizing the characterization of micelles NtL-ButM and Neg-ButM, including hydrodynamic diameter and ζ-potential from DLS, critical micelle concentration, radius of gyration and aggregation number from SAXS.

These block copolymers can be then formulated into nanoscale micelles to achieve high suspensibility in aqueous solutions as well as controlled release of butyrate from the core. The pHPMA-b-pBMA was self-assembled into neutral micelles (NtL-ButM) through a cosolvent evaporation method **(Fig. 1b**). The hydrophobic pBMA block forms the core, while the hydrophilic pHPMA forms the corona. In contrast, pMAA-b-pBMA cannot be formulated into micelles by this method because of the formation of intramolecular hydrogen bonds between pMAA chains^31^. Such bonding can, however, be disrupted when a strong base, here NaOH, is titrated into the mixture of pMAA-b-pBMA polymer to change methacrylic acid into ionized methacrylate^32–34^. Upon base titration, pMAA-b-pBMA polymer can then self-assemble into negatively charged micelles (Neg-ButM) (**Fig. 1b**). Cryogenic electron microscopy (CryoEM) revealed the detailed structure of the micelles, especially the core structure made of pBMA, which was more condensed with higher contrast. CryoEM images indicated that the diameter of the core of NtL-ButM was 30 nm, while Neg-ButM had a smaller core diameter of 15 nm (**Fig. 1c, d**). Both NtL-ButM and Neg-ButM have similar sizes of 44.7 ± 0.8 nm and 39.9 ± 1.6 nm, respectively, measured by dynamic light scattering (DLS) (**Fig. 1e**, **Fig. S10a**). Their low polydispersity index below 0.1 indicated the monodispersity of those micelles. As expected, NtL-ButM has a near-zero ζ-potential of −0.3 ± 0.5 mV, while Neg-ButM’s is −31.5 ± 2.3 mV due to the ionization of methacrylic acid (**Fig. 1e**).To obtain the critical micelle concentration (CMC) of NtL-ButM and Neg-ButM, which indicates the likelihood of formation and dissociation of micelles in aqueous solutions, pyrene was added during the formulation and the fluorescence intensity ratio between the first and third vibronic bands of pyrene was plotted to calculate the CMC (**Fig. S11**)^35^. Results showed that Neg-ButM had a higher CMC of 14.0 ± 3.5 μM, compared to the CMC of NtL-ButM, which was 0.8 ± 0.4 μM (**Fig. 1e**). The higher CMC indicated that Neg-ButM micelles would be easier to dissociate in solution, possibly because the surface charge made the micellar structure less stable compared to the neutral micelle NtL-ButM. In addition, we conducted small angle X-ray scattering (SAXS) analysis on both micelles to obtain the aggregation number (**Fig. S12**). As indicated from Guinier plots, radii of gyration for NtL-ButM and Neg-ButM were 14.2 nm and 13.5 nm (**Fig. 1e**), respectively, and the structures of micelles were confirmed to be spheres from Kratky plots of SAXS data (**Fig. S12**). We then fitted the SAXS data with a polydispersity core-shell sphere model with the assumptions that the micelle has a spherical core with a higher scattering length densities (SLD) and a shell with a lower SLD^36^. The model gave us the volume fraction of the micelles, the radius of the core, and the thickness of the shell, allowing us to calculate the aggregation number and mean distance between micelles. According to the fitting results, aggregation numbers for NtL-ButM and Neg-ButM were 119 and 92, respectively (**Fig. 1e**).

### Butyrate micelles release butyrate in the lower GI tract

Given that butyrate is linked to the micelle-forming chain via ester bonds, we validated the release of butyrate in *ex vivo* conditions, including in simulated gastric fluid and simulated intestinal fluid that mimic those biological environments. In the simulated gastric fluid, both Neg-ButM and NtL-ButM showed negligible release of butyrate within hours, and sustained slow release over three weeks, while Neg-ButM had even slower release rate than NtL-ButM (**Fig. 2a**). The anionic surface of Neg-ButM in the acidic environment is likely responsible for the resistance to hydrolysis of the BMA core. In addition, we observed that the Neg-ButM was not stable in acidic conditions, i.e., in the simulated gastric fluid *in vitro*, and can aggregate into larger polymer particles (**Fig. S13b**). By contrast, in simulated intestinal fluid, both micelles released the most of their butyrate within minutes in the presence of a high concentration of the esterase pancreatin (**Fig. 2b**).

**Fig. 2.**
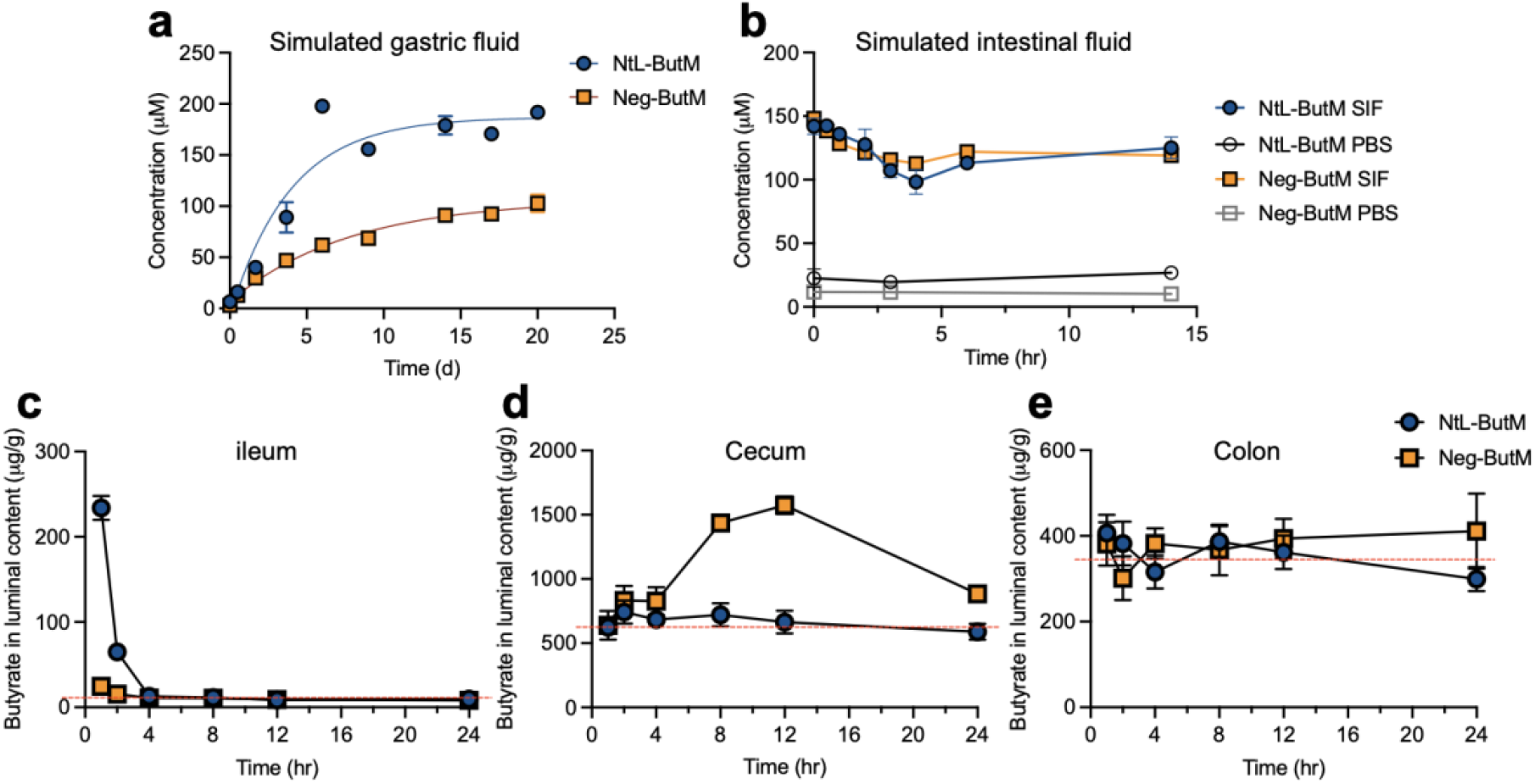
*In vitro* and *in vivo* butyrate release in the GI tract from NtL-ButM and Neg-ButM. **a,** Both NtL-ButM and Neg-ButM released butyrate slowly in the simulated gastric fluid over 20 days. **b,** Both NtL-ButM and Neg-ButM released their complete butyrate load within minutes in simulated intestinal fluid (SIF) containing high levels of the esterase pancreatin. Neither polymer released butyrate in PBS on these timescales. n = 3. **c-e**, The amount of butyrate released in the ileum, cecum, or colon contents after a single intragastric administration of NtL-ButM or Neg-ButM at 0.8 mg/g to SPF C3H/HeJ mice. Butyrate was derivatized with 3-nitrophenylhydrazine and quantified with LC-MS/MS (ileum samples) or LC-UV (cecum and colon samples). The dotted red lines represent butyrate content in untreated mice. n = 9-10 mice per group. Data represent mean ± s.e.m.

We then measured butyrate levels in the mouse GI tract after administering a single dose of NtL-ButM or Neg-ButM by intragastric gavage (i.g.). Both LC-UV and LC-MS/MS methods have been used to measure butyrate concentrations in the luminal contents of the ileum, cecum, and colon, the sites where butyrate producing bacteria normally reside^37,38^. However, because the baseline concentration in the ileum was too low for the UV detector, we used LC-MS/MS to measure the butyrate concentration in that GI tract segment. In SPF (non-antibiotic treated) mice, intragastric administration of NtL-ButM increased the butyrate concentration in the ileum for up to 2 hr after gavage (**Fig. 2c**), but this was short lived, and the butyrate concentration did not increase in either the cecum or colon **(Fig. 2d, e**). Interestingly, Neg-ButM raised butyrate concentrations by 3-fold in the cecum starting from 4 hr after gavage and lasting for at least another 8 hr but not in the ileum or colon (**Fig. 2c-e**). It is possible that the butyrate released in the cecum will continuously flow into the colon; our inability to detect increased concentrations of butyrate in the colon is likely due to its rapid absorption and metabolism by the colonic epithelium. In addition, as expected the polymer backbone of the micelles remained intact when passing through the GI tract. We observed less than 28% molecular weight loss – the percentage of butyrate content – of the polymer in fecal samples collected from 4-8 hr after oral administration (**Fig. S15 a, b**). Moreover, when incubated in a hydrolytic environment *in vitro*, the polymer backbone remained intact after releasing most of the butyrate over 7 days in 125 mM sodium hydroxide solution (**Fig. S15 c, d**).

We further investigated how these two butyrate micelles transit through the GI tract by administering fluorescently labeled NtL-ButM or Neg-ButM to mice i.g. and visualizing their biodistribution via an In Vivo Imaging System (IVIS) (**Fig. S16**). In this case, the fluorescent marker was conjugated to the polymer chain, allowing visualization of the transit of the polymer backbone itself. The IVIS results validated that the polymeric micelles were retained in the mouse GI tract for more than 6 hr after gavage. The neutral micelle NtL-ButM passed through the stomach and small intestine within 2 hr and accumulated in the cecum. However, negatively charged Neg-ButM accumulated in the stomach first and then gradually traveled through the small intestine to the cecum. Overall, Neg-ButM had a longer retention time in the stomach and small intestine, which is possibly due to a stronger adhesive effect to the gut mucosa imparted by the negative charge^39,40^. Both micelles were cleared from the GI tract within 24 hr after administration. In addition, we measured the fluorescence signal from other major organs and plasma by IVIS (**Fig. S16b**), as well as the butyrate concentration in the plasma by LC-MS/MS. The signals were all below the detection limit from both methods, suggesting that there was negligible absorption of these butyrate micelles into the blood circulation from the intestine, consistent with our desire to deliver butyrate to the lower GI tract and to avoid any complexities of systemic absorption of the polymer or micelles.

Due to the existing high level of butyrate produced within the gut in SPF conditions (i.e., with an intact microbiome), we conducted similar biodistribution experiments on vancomycin-treated mice (**Fig. S17a**), where most of the Gram positive bacteria, including the butyrate-producing bacteria, were depleted to induce dysbiosis. We observed that both NtL-ButM and Neg-ButM transited to the lower GI tract within an hour (**Fig. S17b**). Free butyrate from NaBut was measurable in the stomach from 1-4 hours post gavage but was barely detectable in the lower GI sections such as cecum and colon (**Fig S17c**). In contrast, NtL-ButM and Neg-ButM only released 2.3%, or 8.8% of the amount of butyrate in the stomach as compared to NaBut treatment. In antibiotic treated mice, both NtL-ButM and Neg-ButM released butyrate in the cecum and colon. The peak concentrations were observed between 2-4 hours after oral gavage, which is similar to what we observed from fluorescent signals from the IVIS images (**Fig. S17b**).

Delivery of butyrate to the lower GI tract could affect the host immune response by interacting with the intestinal epithelium. To investigate whether and how our butyrate micelles regulate gene expression in the distal small intestine, we performed RNA sequencing of the ileal epithelial cell compartment (**Fig. S18a**). Germ-free (and thus butyrate-depleted) C3H/HeN mice were treated daily with NtL-ButM i.g. for one week and ileal epithelial cells were collected for RNA isolation and sequencing. Because only NtL-ButM (and not Neg-ButM) released butyrate in the ileum, only NtL-ButM was used for this experiment to examine local effects. NtL-ButM-treated mice had unique gene expression signatures compared to those treated with PBS or control polymer, which consists of the same polymeric structure but does not contain butyrate. Interestingly, most genes upregulated by NtL-ButM treatment were Paneth cell-derived antimicrobial peptides (AMPs), including angiogenin 4 (*Ang4*), lysozyme-1 (*Lyz1*), intelectin (*Itln1*) and several defensins (*Defa3, Defa22, Defa24* etc.) (**Fig. S18a, Fig. S19**). We quantified the protein level of intelectin, one of the up-regulated AMPs (**Fig. S18b, c**). We chose intelectin because it is known to be expressed by Paneth cells which reside in small intestinal crypts and can recognize the carbohydrate chains of the bacterial cell wall^41^. Paneth cell AMPs have largely been characterized in C57BL/6 mice and specific reagents are available for their detection in that strain^42^. GF C57BL/6 mice were gavaged daily with NtL-ButM or PBS for one week. Immunofluorescence microscopy of ileal sections revealed that the NtL-ButM treated group expressed a large amount of intelectin in the crypts of the ileal tissue. However, images from the PBS group showed limited intelectin signal (**Fig. S18b**). Quantification using ImageJ of relative fluorescence intensity per ileal crypt also showed that the NtL-ButM group had significantly higher expression of intelectin compared to the PBS control (**Fig. S18c**). The intelectin staining thus further supported the pharmacological effects of NtL-ButM; up-regulation of intelectin induced by NtL-ButM was not only demonstrated on the transcriptional level by RNAseq but was also validated at the protein level. We next performed RT-qPCR on ileal epithelial cells from adult SPF C57BL/6 mice (i.e., mice with a replete butyrate producing microbiota) treated with PBS or NtL-ButM i.g. for one week (**Fig. S20**). In contrast to what we observed in the GF mice, there were no significant differences in the expression of AMPs in SPF mice. It has been shown that these AMPs are constitutively expressed, and very few stimuli abrogate or increase their expression in SPF mice^42,43^. As there is no butyrate deficit in these SPF mice, administration of NtL-ButM did not show the effect of upregulation of AMP gene expression that we observed in GF mice. These results suggest that it is unlikely that ButM acts by modulation of AMPs in mice with an intact microbiota.

### Butyrate micelles repair intestinal barrier function

As discussed above, butyrate-producing bacteria play an important role in the maintenance of the intestinal barrier. To assess the effects of locally delivered butyrate on intestinal barrier integrity, we treated mice with the chemical perturbant DSS for 7 days to induce epithelial barrier dysfunction^44^. Due to the different biodistribution and butyrate release behaviors *in vivo* from the two butyrate micelles, we reasoned that the combined dosing of NtL-ButM and Neg-ButM would cover the longest section of the lower GI tract and last for a longer time; thus, a 1:1 combination of NtL-ButM and Neg-ButM (abbreviated as ButM) was selected for study. Throughout DSS treatment, and for three days after DSS administration was terminated, mice were orally gavaged twice daily with either PBS or ButM at three different concentrations, or once daily with cyclosporin A (CsA) as the positive therapeutic control (as outlined in **Fig. 3a**). Intragastric gavage of 4 kDa FITC-dextran was used to evaluate intestinal barrier permeability. We detected a significantly higher concentration of FITC-dextran in the serum 4 hr after gavage in DSS-treated mice that received only PBS, demonstrating an impaired intestinal barrier. Naïve mice (without DSS exposure), or the DSS-treated mice that also received either CsA or ButM at all three concentrations had similar serum levels of FITC-dextran, suggesting that treatment with ButM successfully repaired the DSS-induced injury to the barrier (**Fig. 3b**). Additionally, neonatal antibiotic treatment impairs homeostatic epithelial barrier function and increases permeability to food antigens^25^. Thus, we further evaluated whether ButM treatment can reduce intestinal barrier permeability in antibiotic-treated mice (**Fig. 3c**). In the antibiotic treatment model, serum was collected 1.5 hr after FITC-dextran gavage, a timepoint at which the luminal FITC-dextran has transited through the ileum but has not yet reached the colon, to specifically analyze barrier permeability in the small intestine. Similar to what we observed in the DSS-induced model, mice treated with ButM had significantly lower FITC-dextran levels in the serum compared to mice that received PBS (**Fig. 3d**), demonstrating that ButM effectively rescued both DSS-induced and antibiotic-induced intestinal barrier dysfunction.

**Fig. 3.**
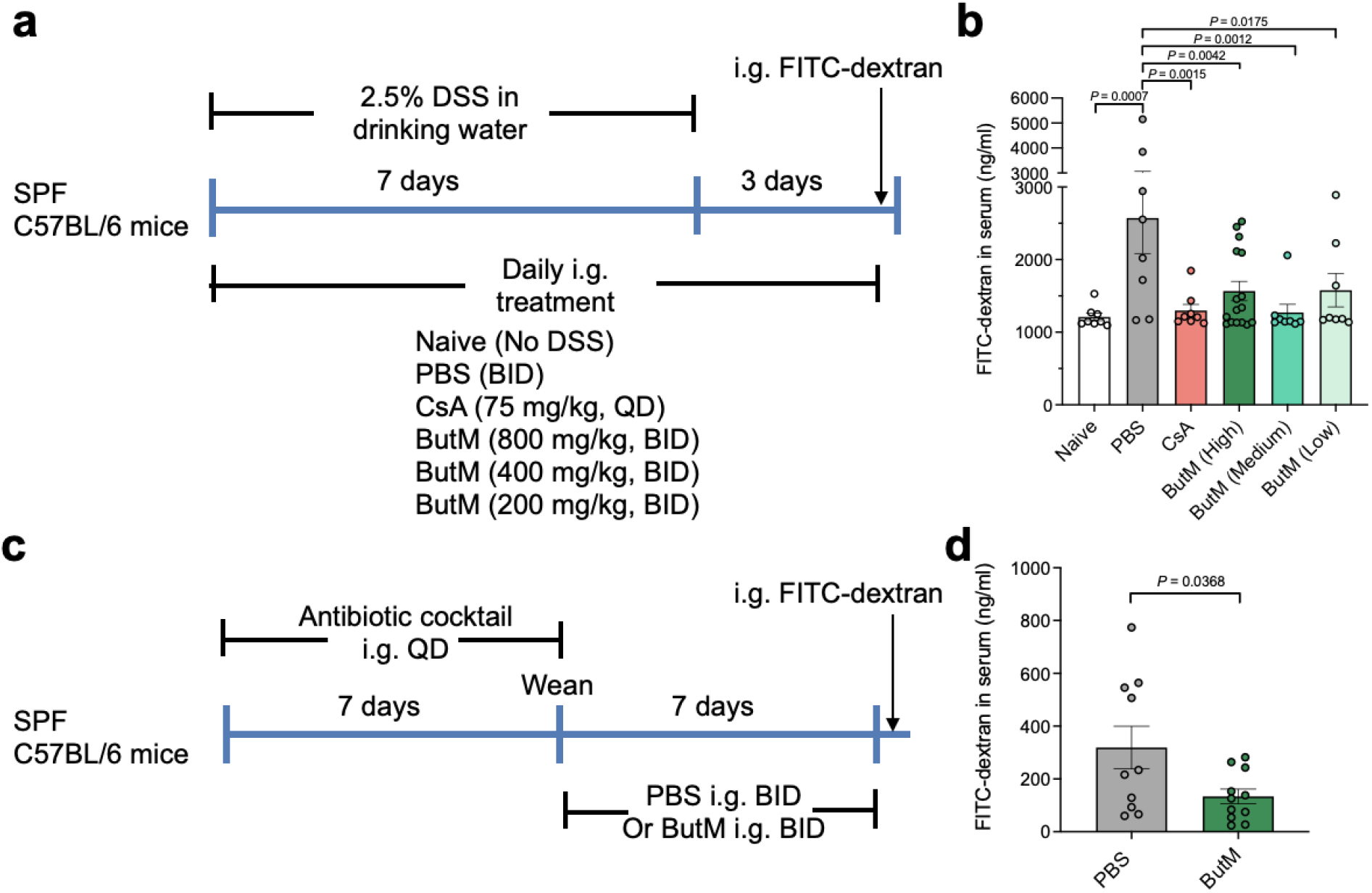
Butyrate micelle treatment repaired intestinal barrier integrity in DSS-treated or antibiotic-treated mice. **a,** Mice were given 2.5% DSS in the drinking water for 7 days to induce epithelial barrier dysfunction. DSS was removed from the drinking water on days 7-10. For treatment mice were intragastrically (i.g.) dosed daily with either PBS, cyclosporin A (CsA), or ButM at different concentrations. QD: once a day, BID: twice daily at 10-12 hr intervals. On day 10, all mice received an i.g. administration of 4kDa FITC-dextran. Fluorescence was measured in the serum 4 hr later. **b,** Concentration of FITC-dextran in the serum. n=8 mice per group, except for high dose ButM which had 16 mice per group. **c**, Mice were treated with a mixture of antibiotics, beginning at 2 wk of age, for 7 days. After weaning, mice were i.g. administered either PBS (n=10) or ButM (n=11) at 800 mg/kg twice daily for 7 days. All mice then received an i.g. administration of 4kDa FITC-dextran. Fluorescence was measured in the serum 1.5 hr later. **d**, Concentration of FITC-dextran in the serum. Data in **d** is pooled from two independent experiments. Data represent mean ± s.e.m. Comparisons were made using one-way ANOVA with Dunnett’s post-test(**c**), or Student’s t-test (**d**).

### Butyrate micelles ameliorated anaphylactic responses in peanut allergic mice

To evaluate the efficacy of the butyrate-containing micelles in treating food allergy, we tested ButM in a well-established murine model of peanut-induced anaphylaxis^25,45^. All of the mice were treated with vancomycin to induce dysbiosis. Beginning at weaning, vancomycin-treated SPF C3H/HeN mice were intragastrically sensitized weekly for 4 weeks with peanut extract (PN) plus the mucosal adjuvant cholera toxin (CT) (**Fig. 4a, b**), as previously described^25,45^. Following sensitization, some of the mice were challenged with intraperitoneal (i.p.) PN and their change in core body temperature was monitored to ensure that the mice were uniformly sensitized; a decrease in core body temperature is indicative of anaphylaxis (**Fig. 4c**). The rest of the sensitized mice were then treated i.g. twice daily for two weeks with either PBS or the combined micelle formulation ButM. After two weeks of therapy, the mice were challenged by i.p. injection of PN and their core body temperature was assessed to evaluate the response to allergen challenge. Compared with PBS-treated mice, allergic mice that were treated with ButM experienced a significantly reduced anaphylactic drop in core body temperature (**Fig. 4d**). In addition, ButM-treated mice also had significantly reduced concentrations of mouse mast cell protease-1 (mMCPT-1), histamine, peanut-specific IgE, and peanut-specific IgG1 detected in the serum (**Fig. 4e-h**). Histamine and mMCPT-1 are released from degranulating mast cells upon allergen cross-linking of IgE and are reliable markers of anaphylaxis. mMCPT-1 is a chmyase expressed by intestinal mucosal mast cells; elevated concentrations of mMCPT-1 increase intestinal barrier permeability during allergic hypersensitivity responses^46,47^. While ButM very effectively reduced the allergic response to peanut exposure, free NaBut had no effect in protecting allergic mice from an anaphylactic response (**Fig. 4i-k**) and did not reduce serum peanut-specific IgE and IgG1 levels (**Fig. 5l, m**). This failure of NaBut to effectively reduce the allergic response to peanut is further evidence for the necessity to deliver butyrate to the lower GI tract. The effects of ButM on the peanut allergic mice were dose-dependent, since we observed that reducing the dose of ButM by half was not as effective as the full dose in protecting mice from an anaphylactic response (**Fig. S21**). We also evaluated and compared each of the two butyrate micelles (NtL-ButM and Neg-ButM) as monotherapies in treating peanut allergy in mice rather than the 1:1 combination (i.e., ButM). Both NtL-ButM and Neg-ButM administered individually significantly reduced the anaphylactic response to peanut challenge, although not as effectively as the ButM polymer combination (**Fig. S21**). Together, these results demonstrate that ButM is highly effective in preventing allergic responses to food in sensitized mice.

**Fig. 4.**
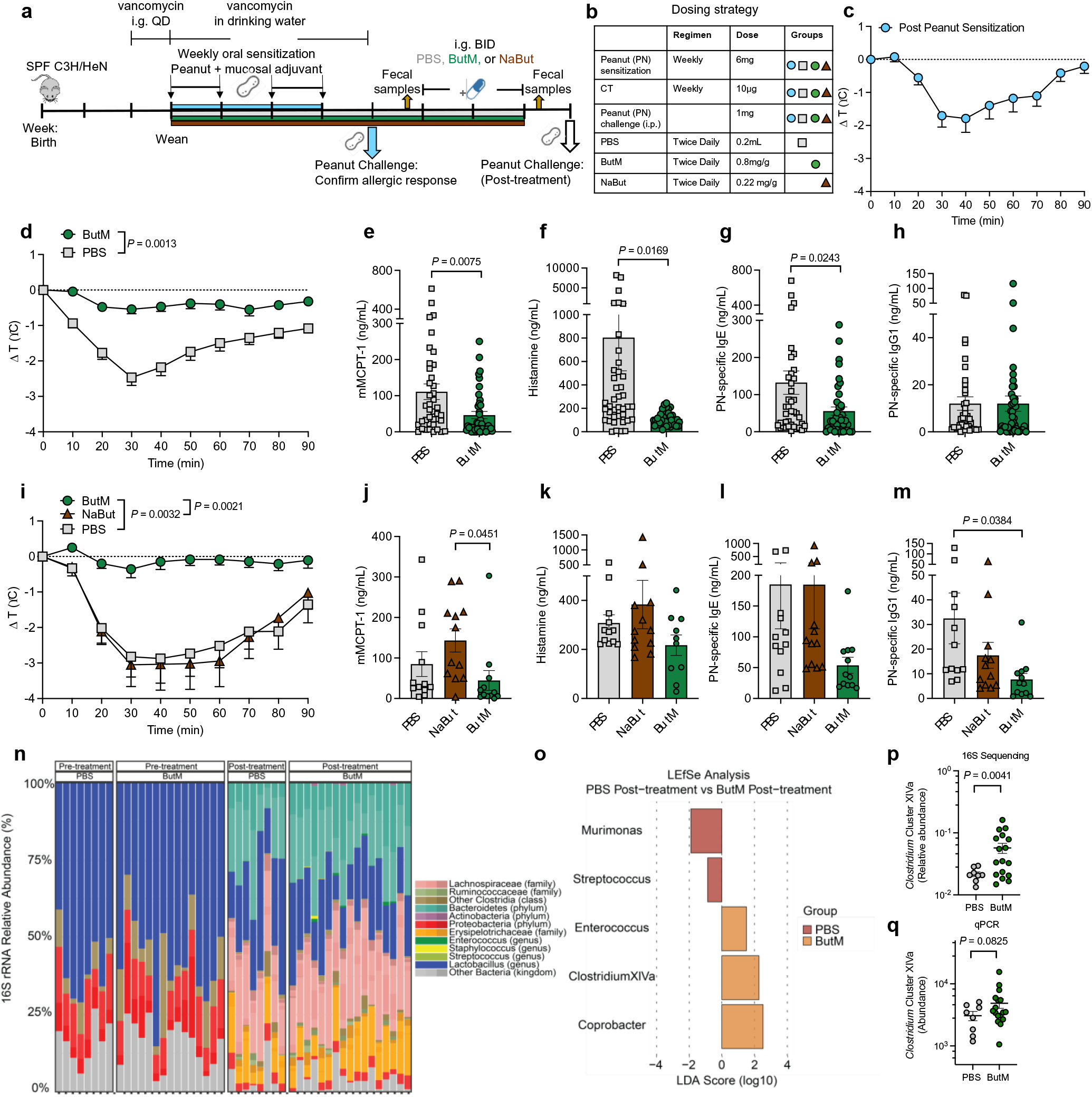
Butyrate micelle treatment reduced the anaphylactic response to peanut challenge and promoted the recovery of *Clostridium* Cluster XIVa after antibiotic exposure. **a, b,** Experimental schema and dosing strategy. All of the mice were sensitized weekly by intragastric gavage of 6 mg of peanut extract (PN) plus 10 μg of the mucosal adjuvant cholera toxin. After 4 weeks of sensitization one group of mice (n=20) was challenged by i.p. administration of 1 mg of PN to confirm that the sensitization protocol induced a uniform allergic response. Fecal samples were collected before and after treatment with ButM for microbiome analysis**. c**, Change in core body temperature following PN challenge where core body temperature drop indicates anaphylaxis. In the first study (**d-h**), the remaining mice were randomized into two treatment groups. One group was treated with PBS (n=40) and the other group was treated with a 1:1 mix of NtL-ButM and Neg-ButM polymers 0.4 mg/g each (ButM, n=40). The data in the **d-h** is pooled from four independent experiments. **d,** Change in core body temperature following challenge with PN in PBS or ButM treated mice. The area under curve (AUC) values were compared between two groups. **e-h,** Serum mMCPT-1 (**e**), histamine (**f**), peanut specific IgE (**g**) and peanut-specific IgG1 (**h**) from mice in **d**. In the second study, mice were treated with PBS (n=12), ButM (n=12), or sodium butyrate (NaBut, n=12) (**i-m**). **i,** Change in core body temperature following challenge with PN in PBS, ButM, or NaBut treated mice. **j-m,** Serum mMCPT-1 (**j**), histamine (**k**), peanut specific IgE (**l**) and peanut-specific IgG1 (**m**) from mice in **i**. **n**, 16S rRNA sequencing analysis of relative abundance of bacterial taxa in fecal samples of allergic mice collected before (left) or after (right) treatment with PBS (n = 8) or ButM (n = 17). **o**, Differentially abundant taxa between mice treated with PBS or ButM after treatment as analyzed by LEfSe. **p-q,** Relative abundance of *Clostridium* Cluster XIVa in fecal samples after treatment with PBS or ButM (from **n**) analyzed from 16S sequencing data (**p**) or analyzed by qPCR (**q**). For **p** and **q**, Student’s t-test with Welch’s correction was used for statistical analysis. Data represent mean ± s.e.m. Data analyzed using two-sided Student’s t-test for **d-h**, and one-way ANOVA with Tukey’s post-test for **i-m**. QD: once a day, BID: twice daily at 10-12 hr intervals.

**Fig. 5.**
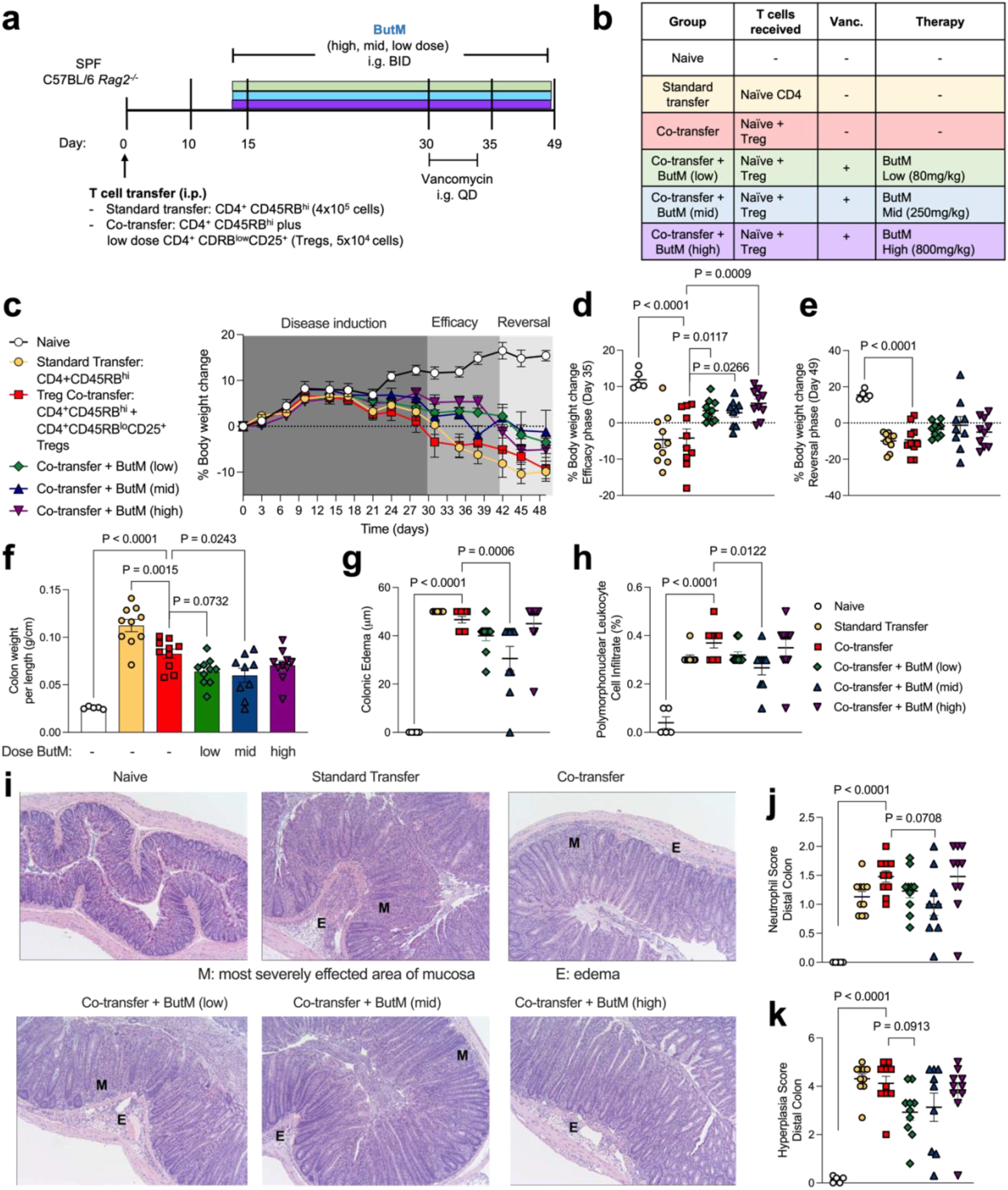
ButM decreases severity of T cell transfer colitis. **a**, Experimental design. On day 0, *Rag2^-/-^* mice received an i.p. injection of purified naïve CD4 T cells (CD4^+^CD45RB^hi^) alone (standard transfer) or together with CD4^+^CD45RB^lo^CD25^+^ Treg cells (co-transfer). Naïve animals received a sham injection. Some groups were gavaged twice daily with ButM at a low (80mg/kg), medium (250mg/kg) or high (800mg/kg) dose from day 14 until end of experiment. ButM treated mice were also gavaged with vancomycin (0.1mg) from day 30-34. **b**, Table detailing treatment groups. **c**, Body weight change as a percent of starting weight over duration of study window. **d**, **e,** Body weight change (% of initial weight) from representative days during the efficacy (**d,** day 35) and reversal (**e,** day 49) phases of the study. **f**, Weight to length ratio of colons harvested on day 49. **g**, Length of edemas from colon histology. **h,** Abundance of polymorphonuclear cell infiltrates (% of cells) from colon histology. **i**, Representative images of transverse sections of distal colon tissue from mice in each treatment group. Tissues were stained with H&E and images were taken at 100x. **j**, Neutrophil score in the distal colon from histological analysis. **k**, Hyperplasia score in the distal colon from histological analysis. Experiment performed by Inotiv (Boulder, CO). In **c**, circles represent mean ± s.e.m. for all mice. In **d-f**, dots represent individual mice, bars represent mean ± s.e.m. In **g, h, j, k,** dots represent average of all histopathological images per mouse, bars represent mean ± s.e.m. across mice. Statistics are analyzed as one-way ANOVA with Dunnett’s multiple comparison’s test comparing all groups against the Treg co-transfer group. QD: once a day, BID: twice daily at 10-12 hr intervals. n = 5 naïve mice, n = 10 for experimental groups.

### Butyrate micelles alter the fecal microbiota and promote recovery of Clostridia after antibiotic exposure

We next examined whether treatment with ButM altered the fecal microbiome. In the mouse model of peanut allergy described above, we induced dysbiosis by treating mice with vancomycin one week before the start of allergen sensitization and throughout the sensitization regimen. Vancomycin depletes Gram positive bacteria, including Clostridial species^48^. After sensitization, we removed vancomycin from the drinking water and compared the fecal microbial composition of the allergic mice before and after treatment with PBS or ButM (see timepoints collected in **Fig. 4a**). 16S rRNA targeted sequencing confirmed depletion of Clostridia in vancomycin-treated mice; the fecal microbiota was instead dominated by Lactobacillus and Proteobacteria (**Fig. 4n**, left). After halting vancomycin administration, regrowth of Clostridia (including Lachnospiraceae and others) and Bacteroidetes was observed in both the PBS and ButM treated groups. This regrowth of Clostridia, and corresponding decrease in relative abundance of Lactobacillus and Proteobacteria, can be observed in the overall taxonomic composition (**Fig. 4n**, right) or by LEfSe analysis comparing pre-and post-treatment timepoints within each treatment group (**Fig S22**). When comparing differentially abundant taxa between treatment groups by LEfSe analysis, *Murimonas* and *Streptococcus* were significantly higher in relative abundance in the PBS post-treatment group when compared to the ButM (**Fig. 4o**). ButM treatment significantly increased the relative abundance of *Enterococcus*, *Coprobacter*, and *Clostridium* Cluster XIVa (**Fig. 4o**). *Clostridium* Cluster XIVa is a numerically predominant group of bacteria (in both mice and humans) that is known to produce butyrate, modulate host immunity, and induce Tregs^49^. We confirmed that the relative abundance of *Clostridium* Cluster XIVa in mice treated with ButM was significantly increased in the 16S data set (**Fig. 4p**) and quantified the enriched abundance of this taxa by qPCR (**Fig. 4q)**. Our finding of increased abundance of *Clostridium* Cluster XIVA after treatment with ButM is in keeping with earlier work which showed that butyrate sensing by peroxisome proliferator-activated receptor (PPAR-γ) shunts colonocyte metabolism toward β-oxidation, creating a local hypoxic niche for these oxygen sensitive anaerobes^50^. We performed a second experiment to examine the effects of ButM treatment on the composition of the microbiota in the absence of allergic sensitization (to rule out a confounding influence of the sensitization protocol itself while holding all other variables constant). Strikingly, we found a similar increase in the relative abundance of Clostridia in the feces and ileal contents of mice treated with ButM (**Fig S23**). In this experiment, the relative abundance of *Clostridium* Cluster IV was increased in the feces, and *Clostridium* Cluster XVIII was increased in both feces and ileal contents.

To gain additional mechanistic insights we also evaluated the effect of ButM on immune cells in the peanut allergy model. We treated peanut-sensitized mice with either PBS, NaBut, or ButM for two weeks and examined the effects of ButM treatment on the abundance of Tregs (**Fig. S24**) and on myeloid cell activation (**Fig. S26**). We found that ButM did not affect the proportion or number of FoxP3^+^CD25^+^ Tregs in the spleen, ileum, or colon-draining lymph nodes (LNs) (**Fig. S24b-g**). However, ButM, but not NaBut, significantly down-regulated the expression MHC Class II and the co-stimulatory marker CD86 on cells in the CD11c^hi^, CD11b^+^F4/80^+^, and CD11b^+^CD11c^-^ compartments from both ileum and colon draining LNs (**Fig. S26)**. This data suggests a potential mechanism of action of ButM through suppressing the myeloid cell activation in the mesenteric LNs.

### Butyrate micelles decrease severity of colitis in the CD45RB^hi^ T cell transfer model

Finally, we examined the efficacy of ButM in the CD45RB^hi^ T cell transfer model. In this model (**Fig. 5a**), adult SPF C57BL/6 *Rag2^-/-^* mice (which lack all lymphocytes) receive intraperitoneal injections of purified CD4^+^ splenic T cells from SPF C57BL/6 mice. The transfer of CD45RB^hi^ effector T cells into a lymphopenic host induces acute inflammation in the colonic tissue and pancolitis^51^. Co-administration of CD45RB^lo^ Tregs prevents the development of colitis. The control group received a standard transfer of only CD4 T effectors (CD4^+^CD45RB^hi^) to demonstrate maximal disease severity. Other experimental groups received co-transfer of CD4^+^CD45RB^hi^ cells plus Tregs (CD4^+^CD25^+^CD45RB^lo^, see table of treatment groups in **Fig. 5b**). Because butyrate can induce Tregs^17–19^, a small number was transferred such that Tregs alone were not sufficient to fully prevent disease progression. Some groups that received the co-transfer of Tregs were then treated, beginning 14 days after adoptive transfer, with one of three doses of ButM (low, medium, high) twice daily for the duration of the experiment. All mice began to develop colitic symptoms around day 21, demonstrated by loss of body weight as percent of initial weight (**Fig. 5c**). Co-transfer of Tregs did not improve body weight compared to standard transfer, as expected for the low number of Tregs transferred. However, treatment with any dose of ButM noticeably improved body weight retention from day 30 to 39 (**Fig. 5c**, efficacy phase). The percent change in body weight on day 35 was significantly improved versus the co-transfer control in all three ButM treatment groups (**Fig. 5d**). Based on the results obtained in **Fig. 4** and **Fig. S23**, we predicted that, after two weeks of twice daily treatment, the efficacy of the ButM micelles would be mediated, at least in part, by an expansion of butyrate producing Clostridia. We therefore administered vancomycin to each of the ButM treatment groups daily from day 30-34 to deplete butyrate producing Clostridia. As predicted, the efficacy of ButM was reduced after vancomycin treatment (reversal phase, **Fig. 5c**) and the percent body weight change in the ButM treated mice (at all doses) was similar to co-transfer controls (**Fig. 5c, e**). However, even after the reversal of efficacy induced by vancomycin, mice treated with the medium dose of ButM had a significantly decreased colon weight-to-length ratio compared to co-transfer controls, indicative of less severe disease (**Fig. 5f**). Histopathological analysis demonstrated that the medium dose of ButM also significantly reduced the size of edema (**Fig. 5g**) and reduced the occurrences of polymorphonuclear leukocyte cell infiltrates (**Fig. 5h**) across the colon. Additionally, the medium dose of ButM modestly decreased the neutrophil score (**Fig. 5j**) and the low dose modestly decreased the hyperplasia score (**Fig. 5k**) in the distal colon. Representative images of the distal colon of each treatment group are shown (**Fig. 5i**), highlighting areas of most severely affected mucosa (**M**) and areas of edema (**E**). These results show that ButM reduces disease severity in the CD45RB^hi^ T cell transfer model of colitis, and that this effect is reversed by treatment with vancomycin.

## Discussion

The prevalence of NCCDs, including food allergy and inflammatory bowel diseases has increased dramatically over the past 20 years, particularly in developed countries^11,52,53^. Lifestyle changes such as reduced consumption of dietary fiber, increased antibiotic use (including in the food chain), and sanitation, have altered populations of commensal microbes. These alterations lead to several negative health effects, including impairment of intestinal barrier function. Modulating the gut microbiome to redirect immunity has become a substantial effort in both academia and industry. However, this has proven difficult: getting selected bacteria, especially obligate anaerobes, to colonize the gut is far from straightforward. Here, we have instead focused on delivering the metabolites that are produced by these bacteria in a more direct manner, since their therapeutic efficacy relies largely on the action of their metabolites.

Specifically, we developed a polymeric nano-scale system to deliver butyrate to localized regions along the GI tract. The system was based on polymeric micelles formed by block copolymers, in which butyrate is conjugated to the hydrophobic block by an ester bond and can be hydrolyzed by esterases in the GI tract for local release. The linked butyrate moieties drive hydrophobicity in that block and, as release occurs, the remainder of the construct (an inert, water-soluble polymer) continues to transit through the lower GI tract until it is excreted. The butyrate-containing block, when forming the core of micelles, was resistant to the acidic environment found in the stomach, which might prevent a burst release there before the micelle’s transit into the intestine. The two butyrate-prodrug micelles, NtL-ButM and Neg-ButM, share similar structures but have corona charges of neutral and negative, respectively. This results in their distinct biodistribution in the lower GI tract, where they can release butyrate in the presence of enzymes. Here, we combined both the neutral and negatively charged micelles to deliver butyrate along the distal gut. We showed that this combined formulation successfully preserves barrier function, reduces severity of colitis, and protects from severe anaphylactic responses with a short-term treatment. In our mouse model of peanut allergy, where the mice were previously exposed to vancomycin to induce dysbiosis, we showed that ButM treatment could favorably increase the relative abundance of protective bacteria, such as *Clostridium* Cluster XIVa. A clinical trial showed that bacteria in *Clostridium* Cluster XIVa may be critical to the success of fecal microbiota transplant for treatment of colitis ^27^. Increasing the abundance of these bacteria may be one mechanism by which ButM treatment improves epithelial barrier function after exposure to DSS or antibiotics or in the pre-clinical models of food allergy or colitis.

We conducted an experiment to further analyze the effects of ButM on immune cells in the peanut allergic mice. When we examined the abundance of Tregs in the mesenteric LNs and spleen, we did not observe any differences among treatment groups, suggesting that ButM did not affect these Treg populations in this model. We further evaluated if ButM had any effect on the myeloid cells in peanut-allergic mice. ButM treatment significantly down-regulated the expression of MHC Class II and the co-stimulatory marker CD86 on the dendritic cells and macrophages in the colon-draining and ileal-draining LNs, suggesting downmodulation of activation and antigen presentation. It has also been previously shown that butyrate inhibits mast cell activation through FcεRI-mediated signaling^54^. This is also consistent with absence of hypothermia and elevated serum mMCPT-1 and histamine (all indicators of anaphylaxis) after peanut exposure in ButM treated mice.

Investigations of the therapeutic potential of butyrate in animal models have supplemented butyrate in the drinking water or diet at a high dose for three or more weeks^15–21^. Such dosing to achieve therapeutic effects from sodium butyrate is challenging for clinical translation, due to the uncontrolled dosing regimen, difficulties to replicate in humans, and the unpleasant odor and taste of butyrate as a sodium salt. Our formulation has incorporated butyrate in the polymeric micelles at a high load (28 wt%) and is able to deliver and release most of the butyrate in the lower GI tract, in a manner that masks butyrate’s taste and smell. Here, we have used a daily dose of 800 mg/kg of total ButM to treat peanut allergic mice for two weeks. This can be translated to ∼65 mg/kg of total ButM (or equivalent butyrate dose of 18.2 mg/kg) human dose given the differences in body surface area between rodents and the human^55^. This butyrate dose in ButM micelles is comparable to other butyrate dosage forms that have been tested clinically^56,57^, however, through the local targeting and sustained release in the lower GI tract, we expect our ButM formulation to achieve higher therapeutic potential in food allergies and beyond.

Our approach is not antigen-specific, since antigen delivery was not part of the treatment regimen: our initial proof-of-concept in peanut allergy could be readily extended to other food allergens, such as other nuts, milk, egg, soy and shellfish. Additionally, this approach may be applicable to inflammatory bowel disease and other diseases caused by hyperinflammation along the GI tract. Moreover, our platform can also be easily adapted to deliver other SCFAs or other microbiome-derived metabolites in a single form or in combination, providing a more controlled and accessible way to achieve potential therapeutic efficacy.

## Materials and Methods

### Materials for polymer synthesis

N-(2-hydroxyethyl) methacrylamide (HPMA) monomer was obtained from Sigma-Aldrich or Polysciences, Inc. Solvents including dichloromethane, methanol, hexanes, and ethanol were ACS reagent grade and were obtained from Fisher Scientific. All other chemicals were obtained from Sigma-Aldrich.

### Synthesis of N-(2-hydroxyethyl) methacrylamide (2)

To synthesize N-(2-hydroxyethyl) methacrylamide (HEMA, **2**), ethanolamine (3.70 mL, 61.4 mmol, 2.0 eq), triethylamine (4.72 mL, 33.8 mmol, 1.1 eq) and 50 mL DCM were added into a 250 mL flask. After the system was cooled by an ice bath, and methacryloyl chloride (**1**, 3.00 mL, 30.7 mmol, 1.0 efq) was added dropwise under the protection of nitrogen. The reaction was allowed to warm to room temperature and reacted overnight. Then the reaction mixture was concentrated by rotary evaporation and purified on a silica column using DCM/MeOH (MeOH fraction v/v from 0% to 5%). The product was obtained as a colorless oil (3.42 g, 86.3%). MS (ESI). C_6_H_11_NO_2_, m/z calculated for [M+H]^+^: 129.08, found: 129.0. ^1^H-NMR (500 MHz, CDCl_3_) δ 1.93 (s, 3H), 3.43 (m, 2H), 3.71 (m, 2H), 5.32 (s, 1H), 5.70 (s, 1H), 6.44 (br s, 1H) (**Fig. S1**).

### Synthesis of N-(2-butanoyloxyethyl) methacrylamide (3)

To synthesize N-(2-butanoyloxyethyl) methacrylamide (BMA, **3**), N-(2-hydroxyethyl) methacrylamide (3.30 mL, 25.6 mmol, 1.0 eq), triethylamine (7.15 mL, 51.2 mmol, 2.0 eq) and 50 mL DCM were added into a 250 mL flask. After the reaction system was cooled by an ice bath, butyric anhydride (5.00 mL, 30.7 mmol, 1.2 eq) was added dropwise under the protection of nitrogen. The system was allowed to react overnight. The reaction mixture was filtered and washed by NH_4_Cl solution, NaHCO_3_ solution, and water. After drying by anhydrous MgSO_4_, the organic layer was concentrated by rotary evaporation and purified on a silica column using DCM/MeOH (MeOH fraction v/v from 0% to 5%). The product was obtained as a pale-yellow oil (4.56 g, 89.6%). MS (ESI). C_10_H_17_NO_3_, m/z calculated for [M+H]^+^: 199.12, found: 199.1. ^1^H-NMR (500 MHz, CDCl_3_) δ 0.95 (t, 3H), 1.66 (m, 2H), 1.97 (s, 3H), 2.32 (t, 2H), 3.59 (dt, 2H), 4.23 (t, 2H), 5.35 (s, 1H), 5.71 (s, 1H), 6.19 (br s, 1H) (**Fig. S2**).

### Synthesis of poly(2-hydroxypropyl methacrylamide) (pHPMA, 5)

pHPMA was prepared using 2-cyano-2-propyl benzodithioate as the RAFT chain transfer agent and 2,2’-Azobis(2-methylpropionitrile) (AIBN) as the initiator. Briefly, HPMA (**4**, 3.0 g, 20.9 mmol, 1.0 eq), 2-cyano-2-propyl benzodithioate (28.3 mg, 0.128 mmol, 1/164 eq), and AIBN (5.25 mg, 0.032 mmol, 1/656 eq) were dissolved in 10 mL MeOH in a 25 mL Schlenk tube. The reaction mixture was subjected to four freeze-pump-thaw cycles. The polymerization was conducted at 70°C for 30 hr. The polymer was precipitated in a large volume of petroleum ether and dried in the vacuum chamber overnight. The product obtained was a light pink solid (1.8 g, 60 %). ^1^H-NMR (500 MHz, DMSO-d6) δ 0.8-1.2 (m, 6H, CH(OH)-CH_3_ and backbone CH_3_), 1.5-1.8 (m, 2H, backbone CH_2_), 2.91 (m, 2H, NH-CH_2_), 3.68 (m, 1H, C(OH)-H), 4.70 (m, 1H, CH-OH), 7.18 (m, 1H, NH) (**Fig. S3**).

### Synthesis of pHPMA-b-pBMA (6)

The block copolymer pHPMA-b-pBMA was prepared using pHPMA (**5**) as the macro-RAFT chain transfer agent and N-(2-butanoyloxyethyl) methacrylamide (**3**) as the monomer of the second RAFT polymerization. Briefly, pHPMA (1.50 g, 0.105 mmol, 1.0 eq), N-(2-butanoyloxyethyl) methacrylamide (4.18 g, 21.0 mmol, 200 eq), and AIBN (8.3 mg, 0.050 mmol, 0.50 eq) were dissolved in 10 mL MeOH in a 50 mL Schlenk tube. The reaction mixture was subjected to four freeze-pump-thaw cycles. The polymerization was conducted at 70°C for 20 hr. The polymer was precipitated in petroleum ether and dried in the vacuum chamber overnight. The product obtained was a light pink solid (4.22 g, 74%). ^1^H-NMR (500 MHz, DMSO-d6) δ 0.80-1.1 (m, 9H, CH(OH)-CH_3_ (HPMA), CH_2_-CH_3_ (BMA), and backbone CH_3_), 1.55 (m, 4H, CH_2_-CH_2_ (BMA) and backbone CH_2_), 2.28 (m, 2H, CO-CH_2_ (BMA)), 2.91 (m, 2H, NH-CH_2_ (HPMA)), 3.16 (m, 2H, NH-CH_2_ (BMA)), 3.67 (m, 1H, CH(OH)-H), 3.98 (m, 2H, O-CH_2_ (BMA)), 4.71 (m, 1H, CH-OH (HPMA)), 7.19 (m, 1H, NH), 7.44 (m, 1H, NH) (**Fig. S4**).

### Synthesis of pMAA (7) and pMAA-b-pBMA (8)

pMAA (**7**) was prepared using 2-cyano-2-propyl benzodithioate as the RAFT chain transfer agent and AIBN as the initiator. Briefly, methacrylic acid (MAA) (4.0 mL, 47.2 mmol, 1.0 eq), 2-cyano-2-propyl benzodithioate (104.4 mg, 0.472 mmol, 1/100 eq), and AIBN (19.4 mg, 0.118 mmol, 1/400 eq) were dissolved in 20 mL MeOH in a 50 mL Schlenk tube. The reaction mixture was subjected to four freeze-pump-thaw cycles. The polymerization was conducted at 70°C for 24 hr. The polymer was precipitated in hexanes and dried in the vacuum oven overnight. The product obtained was a light pink solid (4.0 g, 100 %). ^1^H-NMR (500 MHz, DMSO-d6) δ 0.8-1.2 (m, 3H, backbone CH_3_), 1.5-1.8 (m, 2H, backbone CH_2_), 7.4-7.8 (three peaks, 5H, aromatic H), 12.3 (m, 1H, CO-OH) (**Fig. S5**).

The block copolymer pMAA-b-pBMA (**8**) was prepared using (**7**) pMAA as the macro-RAFT chain transfer agent and (**3**) N-(2-butanoyloxyethyl) methacrylamide (BMA) as the monomer of the second RAFT polymerization. Briefly, pMAA (0.50 g, 0.058 mmol, 1.0 eq), N-(2-butanoyloxyethyl) methacrylamide (1.47 g, 7.38 mmol, 127 eq), and AIBN (2.4 mg, 0.015 mmol, 0.25 eq) were dissolved in 10 mL MeOH in a 25 mL Schlenk tube. The reaction mixture was subjected to four freeze-pump-thaw cycles. The polymerization was conducted at 70°C for 24 hr. The polymer was precipitated in hexanes and dried in the vacuum oven overnight. The product obtained was a light pink solid (1.5 g, 70%). ^1^H-NMR (500 MHz, DMSO-d6) δ 0.8-1.1 (m, 6H, CH_2_-CH_3_ (BMA), and backbone CH_3_), 1.5-1.7 (m, 4H, CH_2_-CH_2_ (BMA) and backbone CH_2_), 2.3 (m, 2H, CO-CH_2_ (BMA)), 3.2 (m, 2H, NH-CH_2_ (BMA)), 4.0 (m, 2H, O-CH_2_ (BMA)), 7.4 (m, 1H, NH), 12.3 (m, 1H, CO-OH) (**Fig. S6**).

### Synthesis of N_3_-PEG_4_-MA (9) and azide-PEG polymer

In order to include an azide group into pHPMA-b-pBMA or pMAA-b-pBMA polymers, monomer N-(2-(2-(2-(2-azidoethoxy)ethoxy)ethoxy)ethyl) methacrylamide (**9**) was synthesized and used in the copolymerization with HPMA or MAA to obtain the hydrophilic block with azide function. Briefly, N_3_-PEG_4_-NH_2_ (0.5 g, 2.14 mmol, 1.0 eq) and triethylamine (0.60 mL, 4.3 mmol, 2.0 eq) were dissolved in anhydrous DCM. After the reaction system was cooled by an ice bath, methacrylic chloride (0.42 mL, 2.6 mmol, 1.2 eq) was added dropwise under the protection of nitrogen. The system was allowed to react overnight. The reaction mixture was filtered and washed by NH_4_Cl solution, NaHCO_3_ solution, and water. After being dried by anhydrous MgSO_4_, the organic layer was concentrated by rotary evaporation and purified on a silica column using DCM/MeOH (MeOH fraction v/v from 0% to 5%). The product obtained was a pale-yellow oil (0.47 g, 73 %). MS (ESI). C_12_H_22_N_4_O_4_, m/z calculated for [M+H]^+^: 287.16, found: 287.2. ^1^H-NMR (500 MHz, CDCl_3_) δ 6.35 (br, 1H), 5.70 (s, 1H), 5.32 (s, 1H), 3.55-3.67 (m, 12H), 3.52 (m, 2H), 3.38 (t, 2H), 1.97 (s, 3H) (**Fig. S7**). Monomer N_3_-PEG_4_-MA was mixed with HPMA or MAA in a 2:98 wt:wt ratio during the RAFT polymerization to obtain N_3_-pHPMA or N_3_-pMAA. Then, the second block of BMA was added to the macro initiator to obtain N_3_-pHPMA-b-pBMA or N_3_-pMAA-b-pBMA, respectively. The synthesis procedures were the same as the previous description.

### Synthesis of N-hexyl methacrylamide (10) and control polymer

To synthesize a control polymer that did not contain butyrate ester, monomer N-hexyl methacrylamide (**11**) was synthesized and used in the polymerization of hydrophobic block. Briefly, hexanamine (5.8 mL, 46.0 mmol, 1.5 eq), triethylamine (4.7 mL, 33.8 mmol, 1.1 eq) and 50 mL DCM were added into a 250 mL flask. After the system was cooled by an ice bath, methacryloyl chloride (3.0 mL, 30.7 mmol, 1.0 eq) was added dropwise under the protection of nitrogen. The reaction was allowed to warm to room temperature and reacted overnight. Then the reaction mixture was concentrated by rotary evaporation and purified on a silica column using DCM/MeOH (MeOH fraction v/v from 0% to 5%). The product obtained was a colorless oil (4.6 g, 88%). MS (ESI). C_11_H_21_NO, m/z calculated for [M+H]^+^: 184.16, found: 184.2. ^1^H-NMR (500 MHz, CDCl_3_) δ 5.75 (br, 1H), 5.66 (s, 1H), 5.30 (s, 1H), 3.31 (t, 2H), 1.96 (s, 3H), 1.54 (m, 2H), 1.28-1.32 (m, 8H), 0.88 (t, 3H) (**Fig. S8**). After the synthesis of pHPMA or pMAA, monomer N-hexyl methacrylamide (**11**) was used in the polymerization of second block instead of N-(2-butanoyloxyethyl) methacrylamide to obtain control polymers as pHPMA-b-pHMA or pMAA-b-pHMA, respectively (**Fig. S9**). The synthesis procedures were the same as described above.

### Formulation of polymeric micelles

NtL-ButM micelle was formulated by cosolvent evaporation method. 80 mg of pHPMA-b-pBMA polymer was dissolved in 10 mL of ethanol under stirring. After the polymer was completely dissolved, the same volume of 1 × PBS was added slowly to the solution. The solution was allowed to evaporate at room temperature for at least 6 hr until ethanol was removed. After the evaporation, the NtL-ButM solution was filtered through a 0.22 μm filter and stored at 4°C. The size of the micelles was measured by DLS.

Neg-ButM micelle was prepared by base titration^32,33^. 60 mg of pMAA-b-pBMA polymer was added to 8 mL of 1 × PBS under vigorous stirring. Sodium hydroxide solution in molar equivalent to methacrylic acid was added to the polymer solution in three portions over the course of 2 hr. After adding base solution, the polymer solution was stirred at room temperature overnight.1 × PBS was then added to reach the target volume and the solution was filtered through a 0.22 μm filter. The pH of the solution was checked to confirm it was neutral, and the size of the micelles was measured by DLS.

### Dynamic light scattering (DLS) characterizations of micelles

DLS data was obtained from a Zetasizer Nano ZS90 (Malvern Instruments). Samples were diluted 400 times in 1 × PBS and 700 μL was transferred to a DLS cuvette for data acquisition. The intensity distributions of DLS were used to determine the hydrodynamic diameter of micelles. For ζ-potential data, micelles were diluted 100 times in 0.1 × PBS (1:10 of 1 × PBS to MilliQ water) and transferred to disposable folded capillary zeta cells for data acquisition.

### Cryogenic electron microscope imaging of micelles

CryoEM images were acquired on a FEI Talos 200kV FEG electron microscope. Polymeric nanoparticle samples were prepared in 1 × PBS and diluted to 2 mg/mL with MilliQ water. 2 μL sample solution was applied to electron microscopy grid (Agar Scientific) with holey carbon film. Sample grids were blotted, and flash vitrified in liquid ethane using an automatic plunge freezing apparatus (Vitrbot) to control humidity (100%) and temperature (20°C). Analysis was performed at −170°C using the Gatan 626 cry-specimen holder (120,000× magnification; −5 μm defocus). Digital images were recorded on an in-line Eagle CCD camera and processed by ImageJ.

### Measurement of critical micelle concentration

The critical micelle concentrations of NtL-ButM and Neg-ButM were determined by a fluorescence spectroscopic method using pyrene as a hydrophobic fluorescent probe^35,58^. A series of polymer solutions with concentration ranging from 1.0 × 10^-4^ to 2.0 mg mL^-1^ were mixed with pyrene solution with a concentration of 1.2 × 10^-3^ mg mL^-1^. The emission spectra of samples were recorded on a fluorescence spectrophotometer (HORIBA Fluorolog-3) at 20°C using 335 nm as excitation wavelength. The ratio between the first (372 nm) and the third (383 nm) vibronic band of pyrene was used to plot against the concentration of the polymer. The data were processed on Prism software and fitted using Sigmoidal model (**Fig. S11**).

### Small angle x-ray scattering analysis of micelles

SAXS samples were made in 1 × PBS and filtered through 0.2 μm filters. All samples were acquired at Stanford Synchrotron Radiation Lightsource, SLAC National Accelerator Laboratory. SAXS data were analyzed by Igor Pro 8 software (**Fig. S12**). To acquire radius of gyration (R_g_), data were plotted as ln(intensity) vs. q^2^ at low q range. Then R_g_ were calculated from the slope of the linear fitting as shown in the equation (1).

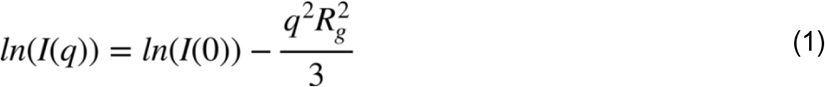

Kratky plot of the data were plotted from I q^2^ vs. q to show the structure of the particles. Moreover, the data were fitted using polydispersed core-shell sphere model (**Fig. S12f, g**)^36^. From the fitting, the radius of the core, thickness of the shell, and volume fraction of the micelle were derived and used to calculate the molecular weight of micelle and the mean distance between micelles using flowing equations:

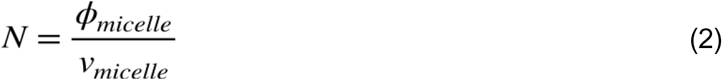

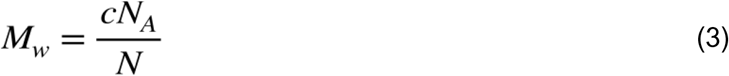

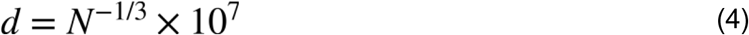

where N is the number of micelles per unit volume. Φ_micelle_ is the volume fraction of micelles derived from fitting. V_micelle_ is the volume of a single micelle, which is calculated from 4/3 πR^3^, where R is the sum of radius of core and thickness of shell. M_w_ is the molecular weight of micelle. C is the polymer concentration. N_A_ is Avogadro constant. D is the mean distance between the micelles in the unit of nm. The aggregation number of micelles were calculated from dividing the molecular weight of micelle by the molecular weight of polymer.

### Mice

C3H/HeN and C3H/HeJ mice were maintained in a Helicobacter, Pasteurella and murine norovirus free, specific pathogen-free (SPF) facility at the University of Chicago. Breeding pairs of C3H/HeJ mice were originally purchased from the Jackson Laboratory. Breeding pairs of C3H/HeN mice were transferred from the germ-free (GF) facility. All experimental mice were bred in house and weaned at 3 weeks of age onto a plant-based mouse chow (Purina Lab Diet 5K67^®^) and autoclaved sterile water. Mice were maintained on a 12 h light/dark cycle at a room temperature of 20–24 °C. GF C3H/HeN or C57BL/6 mice were bred and housed in the Gnotobiotic Research Animal Facility (GRAF) at the University of Chicago. GF mice were maintained in Trexler-style flexible film isolator housing units (Class Biologically Clean) with Ancare polycarbonate mouse cages (catalog number N10HT) and Teklad Pine Shavings (7088; sterilized by autoclave) on a 12 h light/dark cycle at a room temperature of 20–24 °C. All experiments were littermate controlled. All protocols used in this study were approved by the Institutional Animal Care and Use Committee of the University of Chicago. The FITC-dextran intestinal permeability assay in DSS treated mice was performed by Inotiv (Boulder, CO); SPF C57BL/6 mice were obtained from Taconic and housed in the Inotiv animal facility. The T cell transfer colitis model was performed by Inotiv (Boulder, CO) using SPF C57BL/6 donor mice and SPF Ragn12 mice. The studies were conducted in accordance with *The Guide for the Care & Use of Laboratory Animals (8^th^ Edition)* and therefore in accordance with all Inotiv IACUC approved policies and procedures.

### Biodistribution study using in vivo imaging system (IVIS)

SPF C3H/HeJ mice were used for biodistribution studies. Azide labeled pHPMA-b-pBMA or pMAA-b-pBMA polymer was reacted with IR 750-DBCO (Thermo Fisher) and purified by hexane precipitation. After formulation into micelles, the fluorescently labeled NtL-ButM, or Neg-ButM was administered to mice by i.g. gavage. After 1 hr, 3 hr, 6 hr, or 24 hr, mice were euthanized, the major organs were collected from the mice and whole-organ fluorescence was measured via an IVIS Spectrum in vivo imaging system (Perkin Elmer). Images were processed and analyzed by Living Imaging 4.5.5 (Perkin Elmer). In another experiment, SPF C3H/HeJ mice were given 200 mg/L vancomycin in the drinking water for 3 weeks and administered with the same fluorescently labeled NtL-ButM or Neg-ButM by i.g. gavage. Mice were euthanized at 1 hr, 2 hr, 4 hr, 8 hr, 12hr, or 24 hr post-administration, and the whole GI tract was collected for IVIS imaging.

### Butyrate derivatization and quantification using LC-UV or LC-MS/MS

Simulated gastric fluid and simulated intestinal fluid (Fisher Scientific) were used for i*n vitro* release analysis as described previously^59,60^. The simulated gastric fluid was purchased from Ricca Chemical Company, which contains 0.2% (w/v) sodium chloride in 0.7% (v/v) hydrochloric acid and was added with 3.2 mg/mL pepsin from porcine gastric mucosa (Sigma). The simulated intestinal fluid was purchased from Ricca Chemical Company, which contains 0.68% (w/w) potassium dihydrogen phosphate, 0.06% (w/w) sodium hydroxide, and pancreatin at 1% (w/w). NtL-ButM or Neg-ButM were added to simulated gastric fluid or simulated intestinal fluid at a final concentration of 2 mg/mL at 37°C. At pre-determined time points, 20 μL of the solution was transferred into 500 μL of water:acetonitrile 1:1 v/v. The sample was centrifuged using Amicon Ultra filters (Merck, 3 kDa molecular mass cutoff) at 13,000 × g for 15 min to remove polymers. The filtrate was stored at −80°C before derivatization. For the *in vivo* release study in mouse GI tract, NtL-ButM or Neg-ButM micelle solutions were i.g. administered to SPF C3H/HeJ mice at 0.8 mg per g of body weight. Mice were euthanized at 1 hr, 2 hr, 4 hr, 8 hr, 12 hr, and 24 hr after the gavage. Luminal contents from the ileum, cecum, or colon were collected in an EP tube. After adding 500 μL of 1 × PBS, the mixture was vortexed and sonicated for 10 min, and then centrifuged at 13,000 × g for 10 min. The supernatant was transferred and filtered through 0.45 m filter. The filtered solution was stored at −80°C before derivatization.

A similar *in vivo* release experiment was performed in vancomycin-treated SPF C3H/HeJ mice. 4-week-old mice were treated 200 mg/L vancomycin in the drinking water for 3 weeks, followed by oral gavage of sodium butyrate, NtL-ButM or Neg-ButM, at the same butyrate dose of 0.224 mg per g of body weight. At different time points, the stomach, ileum, cecum and colon were harvested. To maximize the butyrate detection, the whole tissue including the content were homogenized in the solution of 1:1 (v/v) water and acetonitrile, and butyrate were extracted in the supernatant after centrifugation and filtered for derivatization and LC-MS/MS measurement.

Sample derivatization for HPLC and LC-MS/MS (**Fig. S14a**): Samples were prepared and derivatized as described in the literature^37^. 3-nitrophenylhydrazine (NPH) stock solution was prepared at 0.02 M in water:acetonitrile 1:1 v/v. EDC stock solution was prepared at 0.25 M in water:acetonitrile 1:1 v/v. 4-methylvaleric acid was added as internal standard. Samples were mixed with NPH stock and EDC stock at 1:1:1 ratio by volume. The mixture was heated by heating block at 60°C for 30 min. Samples were filtered through 0.22 μm filters and transferred into HPLC vials and stored at 4°C before analysis.

LC conditions: The instrument used for quantification of butyrate was Agilent 1290 UHPLC. Column: ThermoScientific C18 4.6 × 50 mm, 1.8 μm particle size, at room temperature. Mobile phase A: water with 0.1% v/v formic acid. Mobile phase B: acetonitrile with 0.1% v/v formic acid. Injection volume: 5.0 μL. Flow rate: 0.5 mL/min. Gradient of solvent: 15% mobile phase B at 0.0 min; 100% mobile phase B at 3.5 min; 100% mobile phase B at 6.0 min; 15% mobile phase B at 6.5 min.

ESI-MS/MS method: The instrument used to detect butyrate was an Agilent 6460 Triple Quad MS-MS. Both derivatized butyrate-NPH and 4-methylvaleric-NPH were detected in negative mode. The MS conditions were optimized on pure butyrate-NPH or 4-methylvaleric-NPH at 1 mM. The fragment voltage was 135 V and collision energy was set to 18 V. Multiple reaction monitoring (MRM) of 222 → 137 was assigned to butyrate (**Fig. S14b**), and MRM of 250 → 137 was assigned to 4-methylvaleric acid as internal standard. The ratio between MRM of butyrate and 4-methylvaleric acid was used to quantify the concentration of butyrate.

### RNA sequencing and data analysis

Starting at the time of weaning, GF C3H/HeN mice were i.g. administered with PBS, NtL-ButM, or control polymer at 0.8 mg/g of body weight once daily for one week. After that time, mice were euthanized, and the ileum tissue was collected and washed thoroughly. The ileal epithelial cells (IECs) were separated from intestinal tissue by inverting ileal tissue in 0.30 mM EDTA, incubating on ice for 30 min with agitation every 5 min. RNA was extracted from the IECs using an RNA isolation kit (Thermo Fisher Scientific) according to manufacturer’s instruction. RNA samples were submitted to the University of Chicago Functional Genomics Core for library preparation and sequencing on a HiSeq2500 instrument (Illumina, Inc.). 50bp single-end (SE) reads were generated. The quality of raw sequencing reads was assessed by FastQC (v0.11.5). Transcript abundance was quantified by Kallisto (v0.45.0) with Gencode gene annotation (release M18, GRCm38.p6), summarized to gene level by tximport (v1.12.3), Trimmed Mean of M-values (TMM) normalized, and log2 transformed. Lowly expressed genes were removed (defined as, counts per million reads mapped [CPM] <3). Differentially expressed genes (DEGs) between groups of interest were detected using limma voom with precision weights (v3.40.6)^61^. Experimental batch and gender were included as covariates for the model fitting. Significance level and fold changes were computed using empirical Bayes moderated t-statistics test implemented in limma. Significant DEGs were filtered by FDR-adjusted *P*<0.05 and fold change ≥ 1.5 or ≤ −1.5. A more stringent *P*-value cutoff (e.g., FDR-adjusted *P*<0.005) may be used for visualization of a select number of genes on expression heatmaps.

### Intelectin stain and microscope imaging

GF C57BL/6 mice were i.g. administered NtL-ButM at 0.8 mg/g of body weight or PBS once daily for one week beginning at weaning. After that time, the mice were euthanized and perfused, small intestine tissue was obtained, rolled into Swiss-rolls, and prepared into tissue section slides. The tissue section slides were fixed and stained with a rat anti-intelectin monoclonal antibody (R&D Systems, Clone 746420) with fluorescent detection and DAPI (ProLong antifade reagent with DAPI). The slides were imaged using a Leica fluorescence microscope. Images were processed by ImageJ software and data were plotted and analyzed by Prism software.

### RT-qPCR of epithelial cells

Intestinal epithelial cells (IECs) were isolated from underlying lamina propria by incubating in 0.03 M EDTA plus 0.0015 M DTT on ice for 20 minutes, then in 0.03 M EDTA for 10 minutes at 37°C. IECs were re-suspended in Trizol® and stored in −80°C until RNA extraction. For Paneth cell product expression, RNA was isolated from homogenized tissue using a guanidine thiocyanate – cesium chloride gradient method, previously described in detail^42,62–64^. cDNA synthesis was performed using the Superscript III reverse transcriptase kit (Invitrogen, Carlsbad, CA), as outlined by the manufacturer. The cDNA was purified using a Qiagen PCR purification kit (Qiagen, Valencia, CA), and then diluted at a concentration representing 10 ng/µl of total RNA input per reaction. The RT-qPCR reactions were performed using a Roche Diagnostics Lightcycler 2.0 (Roche, Indianapolis, IN), using a protocol previously described in detail^42,63^. Primers used to amplify beta actin (*Actb*), α-defensins (*Defa3*, *Defa5*, *Defa20*, *Defa21*, *Defa22*, *Defa23*, *Defa24*, and *Defa26*), lysozyme (*Lyz1*), intelectin-1 (*Itln1*) and solute carrier family 10 member 2 (*Slc10a2*) were reported previously, and are summarized in supplemental Table S1. The quantitative assays utilized to enumerate absolute transcript copy number were as reported^42,63–64^.

### *In vivo* FITC-dextran permeability assay

SPF C57BL/6 8-10 wks old female mice were treated with 2.5% DSS in their drinking water for 7 days. The mice received intragastric administration twice daily, at approximately 10-12 hr intervals, of either PBS or ButM (800, 400 or 200 mg/kg), or once daily with CsA at 75 mg/kg as the positive treatment control. On day 7, DSS was removed from the drinking water for the remainder of the study. On day 10, mice were fasted for 3 hr and dosed with 0.1 mL of FITC-dextran 4kDa (at 100 mg/mL). 4 hr post dose mice were anesthetized with isoflurane and bled to exsanguination followed by cervical dislocation. The concentration of FITC in the serum was determined by spectrofluorometry using as standard serially diluted FITC-dextran. Serum from mice not administered FITC-dextran was used to determine the background. A similar permeability assay was also performed in the antibiotic-depletion model as previously described^25^. Littermate-controlled SPF C57BL/6 mice at 2 wks of age were gavaged daily with a mixture of antibiotics (0.4 mg kanamycin sulfate, 0.035 mg gentamycin sulfate, 850U colistin sulfate, 0.215 mg metronidazole, and 0.045 mg vancomycin hydrochloride in 100 μL PBS) for 7 days until weaning. At weaning, mice were then treated with either PBS or ButM (0.8mg/g) twice daily for 7 days. After the final treatment, the mice were fasted for 3 hr and dosed with 50mg/kg body weight of FITC-dextran 4kDa (at 50 mg/mL). Blood was collected at 1.5 hr post-administration via cheek bleed and the concentration of FITC in the serum was measured as described above.

### CD45RB^hi^ T cell transfer model of colitis

CD4^+^CD45RB^hi^ and CD4^+^CD45RB^low^CD25^+^ cells were isolated from splenocytes of SPF C57BL/6 mice using the CD4 Cell Enrichment Kit (STEMCELL), followed by cell sorting after staining with APC anti-CD4 (clone GK1.5, BioLegend), FITC anti-CD45RB (clone MB4B4, BioLegend), and PE anti-CD25 (clone PC61, BioLegend) antibodies^51^. 6-7 week old female SPF Rag12n mice were injected i.p. with either 4×10^5^ CD45RB^hi^ cells (standard transfer), or with a co-transfer of CD45RB^hi^ cells (minimum 4×10^5^) and CD4^+^CD45RB^low^CD25^+^ Tregs (5×10^4^) in a volume of 200 μl. On the day of transfer, mice were randomly sorted into treatment groups by body weight. Body weight was measured approximately every three days for the duration of the experiment to examine incidence of colitis. Beginning two weeks after T cell transfer, some groups received twice daily intragastric gavages of ButM at one of three doses (low, 80mg/kg; med, 250mg/kg; high, 800mg/kg) for the duration of the experiment. Beginning on day 30 after T cell transfer, mice receiving ButM treatment additionally received a daily gavage of vancomycin (1mg/ 100 μl) for 5 days. Controls received water gavage. Mice were euthanized on day 50 after T cell transfer, and at necropsy weight and length of colon tissue was measured. Tissue sections from the proximal and distal colon were stored in 10% neutral buffered formalin (NBF) for histological analysis.

### Colon histology

Proximal and distal colon tissues of mice from the CD45RB^hi^ T cell transfer colitis model were paraffin embedded, cut in transverse sections, and stained with hematoxylin & eosin (H&E). Edema was quantified as the distance from the outer muscle layer to the muscularis mucosa. Tissue hyperplasia is quantified in a scoring system determined from the size of hyperplasia region (0 = normal, <200μm; 0.5 = very minimal, 201-250μm; 1 = minimal, 251-350μm; 2 = mild, 351-450μm; 3 = moderate, 451-550μm; 4 = marked, 551-650μm; 5 = severe, >650μm). The total inflammation is evaluated in a scoring system determined by extent of immune cell infiltration. The approximate percent of polymorphonuclear leukocytes (PMN) thought to be neutrophils are presented as PMN %. The PMN score is then multiplied by the overall inflammation score to generate a neutrophil score. Histological quantifications are either presented for individual colonic region (distal) or averaged across a distal and proximal region per mouse (whole colon).

### Peanut sensitization, ButM treatment and challenge

SPF C3H/HeN mice were treated with 0.45 mg of vancomycin in 0.1 mL by intragastric gavage for 7 days pre-weaning and then with 200 mg/L vancomycin in the drinking water throughout the remainder of the sensitization protocol. Age- and sex-matched 3-wk-old littermates were sensitized weekly by intragastric gavage with defatted, in-house made peanut extract prepared from unsalted roasted peanuts (Hampton Farms, Severn, NC) and cholera toxin (CT) (List Biologicals, Campbell, CA) as previously described^25,45^. Sensitization began at weaning and continued for 4 weeks. Prior to each sensitization the mice were fasted for 4-5 hr and then given 200 μl of 0.2M sodium bicarbonate to neutralize stomach acids. 30 min later the mice received 6 mg of peanut extract and 10 μg of cholera toxin (CT) in 150 μl of PBS by intragastric gavage. After 4 weeks of sensitization, mice were permitted to rest for 1 wk before a subset of mice was challenged by intraperitoneal (i.p.) administration of 1 mg peanut extract in 200 μl of PBS to confirm that the sensitization protocol induced a uniform allergic response. Rectal temperature was measured immediately following challenge every 10 minutes for up to 90 min using an intrarectal probe, and the change in core body temperature of each mouse was recorded. The remaining mice were not challenged and were randomly assigned into experimental groups. In **Fig. 4** one group of mice was treated with ButM twice daily by intragastric gavage at 0.8 mg of total polymer per gram of mouse body weight (0.8 mg/g) for two weeks, one group of mice received PBS, and another group of mice received sodium butyrate twice daily at 0.224 mg/g (equivalent amount of butyrate to 0.8 mg/g ButM). In the dose-dependent study (**Fig. S21**), mice were treated with either PBS, ButM at 0.8 mg/g (full dose) or ButM at 0.4 mg/g (half dose) twice daily. In addition, groups of mice received NtL-ButM or Neg-ButM only at the full dose, twice daily. After the treatment window, mice were challenged with i.p. administration of 1 mg peanut extract and core body temperature was measured for 90 min. Serum was collected from mice 90 minutes after challenge for measurement of mMCPT-1 and histamine and additionally at 24 hr after challenge for measurement of peanut-specific IgE and IgG1. Collected blood was incubated at room temperature for 1 hour and centrifuged for 7 minutes at 12,000 g at room temperature, and sera were collected and stored at ^-^80°C before analysis. Serum antibodies and mMCPT-1 were measured by ELISA.

### Measurement of mouse mast cell protease 1 (mMCPT-1), histamine, and serum peanut-specific IgE and IgG1 antibodies using ELISA

mMCPT-1 was detected using the MCPT-1 mouse uncoated ELISA kit (ThermoFisher) following the protocol provided by manufacturer. Histamine was assayed using Histamine EIA Kit (Oxford Biomedical Research, Oxford, MI) according to the manufacturer’s protocol. For the peanut specific IgE ELISA, sera from individual mice were added to peanut coated Maxisorp Immunoplates (Nalge Nunc International, Naperville, IL). Peanut-specific IgE Abs were detected with goat anti-mouse IgE-unlabeled (Southern Biotechnology Associates, Birmingham, AL) and rabbit anti-goat IgG-alkaline phosphatase (Invitrogen, Eugene, Oregon) and developed with p-nitrophenyl phosphate “PNPP” (SeraCare Life Sciences, Inc. Milford, MA). For peanut-specific IgG1 ELISA, sera from individual mice were added to peanut coated Maxisorp Immunoplates (Nalge Nunc International, Naperville, IL). Peanut-specific IgG1 was detected using goat anti-mouse IgG1-HRP conjugated (Southern Biotechnology Associates) and substrate TMB liquid substrate system for ELISA (Sigma-Aldrich, St. Louis, MO). The plates were read in an ELISA plate reader at 405 nM (IgE) or 450 nm (IgG1). OD values were converted to nanograms per milliliter of IgE or IgG1 by comparison with standard curves of purified IgE or IgG1 by linear regression analysis and are expressed as the mean concentration for each group of mice ± s.e.m.

### Flow cytometric analysis on immune cells isolated from spleen and mesenteric LNs

Peanut sensitized mice were intragastrically administered twice daily with PBS, sodium butyrate, or ButM at an equivalent dose of 0.224 mg/g (butyrate/mice) for two weeks. After the treatment, spleen, ileum and colon draining LNs from mice were collected, and digested in DMEM supplemented with 5% FBS, 2.0 mg/mL collagenase D (Sigma Aldrich) and 1.2 mL CaCl_2._ Single-cell suspensions were prepared by mechanically disrupting the tissues through a cell strainer (70 μm, Thermo Fisher). Splenocytes (4 x 10^6^) or cells from LNs (1 x 10^6^) were plated in a 96 well-plate. Cells were stained with LIVE/DEAD™ Fixable Aqua Dead Cell Stain Kit (Thermo Fisher), followed by surfacing staining with antibodies in PBS with 2% FBS, and intracellular staining according to the manufacturer’s protocols from eBioscience™ Foxp3/Transcription Factor Staining Buffer Set (Invitrogen). The following anti-mouse antibodies were used: CD3 APC/Cy7 (clone 145-2C11, BD Biosciences), CD4 BV605 (clone RM4-5, Biolegend), CD25 PE/Cy7 (clone PC61, Biolegend), Foxp3 AF488 (clone MF23, BD Biosciences), CD11b BV711 (clone M1/70, BD Biosciences), CD11c PE/Cy7 (clone HL3, BD Biosciences), F4/80 APC (clone RM8, Biolegend), I-A/I-E (MHCII) APC/Cy7 (clone M5/114.15.2, Biolegend), and CD86 BV421 (clone GL-1, Biolegend). Stained cells were analyzed using an LSR Fortessa flow cytometer (BD Biosciences).

### 16S rRNA targeted sequencing

Bacterial DNA was extracted using the QIAamp PowerFecal Pro DNA kit (Qiagen). The V4-V5 hypervariable region of the 16S rRNA gene from the purified DNA was amplified using universal bacterial primers – 563F (5’-nnnnnnnn-NNNNNNNNNNNN-AYTGGGYDTAAA-GNG-3’) and 926R (5’-nnnnnnnn-NNNNNNNNNNNN-CCGTCAATTYHT-TTRAGT-3’), where ‘N’ represents the barcodes, ‘n’ are additional nucleotides added to offset primer sequencing. Illumina sequencing-compatible Unique Dual Index (UDI) adapters were ligated onto pools using the QIAsep 1-step amplicon library kit (Qiagen). Library QC was performed using Qubit and Tapestation before sequencing on an Illumina MiSeq platform at the Duchossois Family Institute Microbiome Metagenomics Facility at the University of Chicago. This platform generates forward and reverse reads of 250 bp which were analyzed for amplicon sequence variants (ASVs) using the Divisive Amplicon Denoising Algorithm (DADA2 v1.14)^66^. Taxonomy was assigned to the resulting ASVs using the Ribosomal Database Project (RDP) database with a minimum bootstrap score of 50^67^. The ASV tables, taxonomic classification, and sample metadata were compiled using the phyloseq data structure^68^. Subsequent 16S rRNA relative abundance analyses and visualizations were performed using R version 4.1.1 (R Development Core Team, Vienna, Austria).

### Microbiome analysis

To identify changes in the microbiome across conditions, a linear discriminant analysis effect size (LEfSe) analysis was performed in R using the microbiomeMarker package and the run_lefse function^69,70^. Features, specifically taxa, can be associated with or without a given condition (e.g., ButM post-treatment vs PBS post-treatment) and an effect size can be ascribed to that difference in taxa at a selected taxonomic level (LDA score). For the LefSe analysis, genera were compared as the main group, a significance level of 0.05 was chosen for both the Kruskall-Wallis and Wilcoxon tests and a linear discriminant analysis cutoff of 1.0 was implemented. The abundance of *Clostridium* Cluster XIVA in post-treatment samples was also determined by quantitative PCR (qPCR) using the same DNA analyzed by 16S rRNA targeted sequencing. Commonly used primers 8F^71^ and 338R^72^ were used to quantify total copies of the 16S rRNA gene for normalization purposes. Primers specific for *Clostridium* Cluster XIVa^73^ were validated by PCR and qPCR. Primer sequences are listed in **Supplementary Table S1**. qPCR was performed using PowerUp SYBR green master mix (Applied Bioystems) according to manufacturer’s instructions. The abundance of *Clostridium* Cluster XIVa is calculated by 2^-CT^, multiplied by a constant to bring all values above 1 (1 x 10^16^), and expressed as a ratio to total copies 16S per gram of raw fecal content.

### Statistical analysis

Statistical analysis and plotting of data were performed using Prism 9.0 (Graphpad), as indicated in the figure legends. One-way ANOVA with Dunnett’s or Tukey’s post-test for multiple comparisons were used in **Fig. 3b, Fig. 4j-m**, and **Fig. 5d-h, j-k**. Two-sided Student’s t-test was used in **Fig. 3d** and **Fig. 4d-h, p-q**. In **Fig. 4d, i**, the area under curve (AUC) values of temperature changes were compared using two-sided Student’s t-test (i) or one-way ANOVA with Tukey’s post-test (d). Data represent mean ± s.e.m.; *n* is stated in the figure legend.

### Code availability

Custom codes and scripts on the RNA sequencing analysis, genomic analysis, and microbiome analysis are available from the corresponding author upon request.

### Data availability

The data that support the findings of this study are available from the corresponding authors upon request. The 16S rRNA and RNAseq raw FastQ data files have been deposited into the National Center for Biotechnology (NCBI) Sequence Read Archive are available under accession numbers PRJNA863725 and (to follow), respectively.

## Acknowledgements

This work was supported by a sponsored research agreement from ClostraBio, Inc., and partially supported by the Chicago Immunoengineering Innovation Center of the University of Chicago. D.S.W. was supported by a fellowship from the Whitaker Foundation. We thank Altayeb A. Alshaikh, Zahra Khosravi, Ha-Na Shim, Nidhi Talasani, Hao Wu, Matthew Bauer and Suzana Gomes for technical assistance. We thank Dr. Betty Theriault from GRAF and Pieter Faber from the Functional Genomics Facility. Parts of this work were carried out at the Cytometry and Antibody Technology Core Facility (Cancer Center Support Grant P30CA014599), the Duchossois Family Institute Microbiome Metagenomics Facility, the Soft Matter Characterization Facility, the Mass Spectrometry Facility (NSF instrumentation grant CHE-1048528), the Nuclear Magnetic Resonance Facility, the Advanced Electron Microscopy Facility (RRID:SCR_019198), and the Functional Genomics Core at the University of Chicago, and the SLAC National Accelerator Laboratory. Some of the bioinformatics analysis was performed on the high-performance computing (HPC) clusters at the University of Chicago Center for Research Informatics, and we thank M. Jarsulic for his technical assistance on the HPC clusters.

## Author contributions

C.R.N. and J.A.H. oversaw all research. R.W., S.C., M.E.H.B., L.A.H., C.R.N., and J.A.H. designed the research strategy. R.W., D.S.W, and J.A.H. conceptualized materials. R.W. and S.C. synthesized materials and fabricated micelles. R.W., S.C., M.E.H.B, L.A.H., Y.S., S.M.C.H., E.C. M.S., and E.B.N. performed experiments. R.W., S.C., M.E.H.B, C.L.B, and L.A.H. analyzed experiments. R.B. performed data analysis on the RNA sequencing experiment, N.P.D. and E.C. performed 16S rRNA targeted microbiome analysis. R.W., S.C., M.E.H.B. L.A.H., C.R.N, and J.A.H. wrote the manuscript. All authors contributed to the article and approved the submitted version.

## Competing interests

C.R.N. and J.A.H. are founders and shareholders in ClostraBio, Inc, which is developing the technology described herein. R.W., S.C., M.E.H.B., D.S.W., C.R.N. and J.A.H. are inventors on patents filed by the University of Chicago describing the micelles reported herein.

## Notes

### Summary of Updates

Figure 5 revised.

## References

1. Iweala, O.I. & Nagler, C.R. The Microbiome and Food Allergy. Annual Review of Immunology 37, 377–403 (2019).

2. Honda, K. & Littman, D.R. The microbiota in adaptive immune homeostasis and disease. Nature 535, 75–84 (2016).

3. Belkaid, Y. & Harrison, O.J. Homeostatic Immunity and the Microbiota. Immunity 46, 562–576 (2017).

4. Wells, J.M., et al. Homeostasis of the gut barrier and potential biomarkers. Am J Physiol Gastrointest Liver Physiol 312, G171–g193 (2017).

5. Donohoe, D.R., et al. The microbiome and butyrate regulate energy metabolism and autophagy in the mammalian colon. Cell metabolism 13, 517–526 (2011).

6. Koh, A., De Vadder, F., Kovatcheva-Datchary, P. & Bäckhed, F. From Dietary Fiber to Host Physiology: Short-Chain Fatty Acids as Key Bacterial Metabolites. Cell 165, 1332–1345 (2016).

7. Berni Canani, R., et al. Lactobacillus rhamnosus GG-supplemented formula expands butryate-producing bacterial strains in food allergic infants. ISMEJ 10, 742–40 (2016

8. Feehley, T., et al. Healthy infants harbor intestinal bacteria that protect against food allergy. Nature Medicine 25, 448–453 (2019).

9. Cait, A. et al. Reduced genetic potential for butyrate fermentation in the gut microbiome of infants who develop allergic sensitization. J. Allergy. Clin Immunol. 144, 1638–1647 (2019).

10. Bao, R., et al. Fecal microbiome and metabolome differ in healthy and food-allergic twins. J Clin Invest 131, e141935 (2021).

11. Lee, M. & Chang, E.B. Inflammatory Bowel Diseases (IBD) and the Microbiome-Searching the Crime Scene for Clues. Gastroenterology 160, 524–537 (2021).

12. Wang W. et al. Increased proportions of Bifidobacterium and the Lactobacillus group and loss of butyrate-producing bacteria in inflammatory bowel disease. J Clin Microbiol. 52, 398–406 (2014)

13. Machiels, K., et al. A decrease of the butyrate-producing species *Roseburia hominis*; and *Faecalibacterium prausnitzii*; defines dysbiosis in patients with ulcerative colitis. Gut 63, 1275–1283 (2014).

14. Liu, H., et al. Butyrate: A Double-Edged Sword for Health? Adv Nutr 9, 21–29 (2018).

15. Tan, J., et al. Dietary Fiber and Bacterial SCFA Enhance Oral Tolerance and Protect against Food Allergy through Diverse Cellular Pathways. Cell Reports 15, 2809–2824 (2016).

16. Sun, M., et al. Microbiota-derived short-chain fatty acids promote Th1 cell IL-10 production to maintain intestinal homeostasis. Nature Communications 9, 3555 (2018).

17. Furusawa, Y., et al. Commensal microbe-derived butyrate induces the differentiation of colonic regulatory T cells. Nature 504, 446–450 (2013).

18. Smith, P.M., et al. The microbial metabolites, short-chain fatty acids, regulate colonic Treg cell homeostasis. Science 341, 569–573 (2013).

19. Arpaia, N., et al. Metabolites produced by commensal bacteria promote peripheral regulatory T-cell generation. Nature 504, 451–455 (2013).

20. Cait, A., et al. Microbiome-driven allergic lung inflammation is ameliorated by short-chain fatty acids. Mucosal Immunology 11, 785–795 (2018).

21. Chen, G., et al. Sodium Butyrate Inhibits Inflammation and Maintains Epithelium Barrier Integrity in a TNBS-induced Inflammatory Bowel Disease Mice Model. EBioMedicine 30, 317–325 (2018).

22. Luceri, C., et al. Effect of butyrate enemas on gene expression profiles and endoscopic/histopathological scores of diverted colorectal mucosa: A randomized trial. Dig Liver Dis 48, 27–33 (2016).

23. Kelly, C.J., et al. Crosstalk between Microbiota-Derived Short-Chain Fatty Acids and Intestinal Epithelial HIF Augments Tissue Barrier Function. Cell Host Microbe 17, 662–671 (2015).

24. Tan, J., et al. The role of short-chain fatty acids in health and disease. Adv Immunol 121, 91–119 (2014).

25. Stefka, A.T., et al. Commensal bacteria protect against food allergen sensitization. Proc Natl Acad Sci U S A 111, 13145–13150 (2014).

26. Halfvarson, J., et al. Dynamics of the human gut microbiome in inflammatory bowel disease. Nature Microbiology 2, 17004 (2017).

27. Wei, S., et al. Gut microbiota differs between treatment outcomes early after fecal microbiota transplantation against recurrent Clostridioides difficile infection. Gut Microbes 14, 2084306 (2022).

28. Rossen, N.G., et al. Findings From a Randomized Controlled Trial of Fecal Transplantation for Patients With Ulcerative Colitis. Gastroenterology 149, 110–118.e114 (2015).

29. Nagler, C.R. Drugging the microbiome. J Exp Med 217, e20191642 (2020).

30. Jimenez, M., Langer, R. & Traverso, G. Microbial therapeutics: New opportunities for drug delivery. J Exp Med 216, 1005–1009 (2019).

31. Xu, F., Xu, J.W. & Luo, Y.L. Impact of hydrogenation on physicochemical and biomedical properties of pH-sensitive PMAA-b-HTPB-b-PMAA triblock copolymer drug carriers. J Biomater Appl 30, 1473–1484 (2016).

32. Colombani, O., et al. Synthesis of Poly(n-butyl acrylate)-block-poly(acrylic acid) Diblock Copolymers by ATRP and Their Micellization in Water. Macromolecules 40, 4338–4350 (2007).

33. Colombani, O., et al. Structure of Micelles of Poly(n-butyl acrylate)-block-poly(acrylic acid) Diblock Copolymers in Aqueous Solution. Macromolecules 40, 4351–4362 (2007).

34. Felber, A.E., Dufresne, M.-H. & Leroux, J.-C. pH-sensitive vesicles, polymeric micelles, and nanospheres prepared with polycarboxylates. Advanced Drug Delivery Reviews 64, 979–992 (2012).

35. Aguiar, J., Carpena, P., Molina-Bolívar, J.A. & Carnero Ruiz, C. On the determination of the critical micelle concentration by the pyrene 1:3 ratio method. Journal of Colloid and Interface Science 258, 116–122 (2003).

36. Wu, H., Ting, J.M., Weiss, T.M. & Tirrell, M.V. Interparticle Interactions in Dilute Solutions of Polyelectrolyte Complex Micelles. ACS Macro Letters 8, 819–825 (2019).

37. Torii, T., et al. Measurement of short-chain fatty acids in human faeces using high-performance liquid chromatography: specimen stability. Ann Clin Biochem 47, 447–452 (2010).

38. Tyagi, A.M., et al. The Microbial Metabolite Butyrate Stimulates Bone Formation via T Regulatory Cell-Mediated Regulation of WNT10B Expression. Immunity 49, 1116–1131.e1117 (2018).

39. Lele, B.S. & Hoffman, A.S. Mucoadhesive drug carriers based on complexes of poly(acrylic acid) and PEGylated drugs having hydrolysable PEG-anhydride-drug linkages. J Control Release 69, 237–248 (2000).

40. Serra, L., Doménech, J. & Peppas, N.A. Engineering design and molecular dynamics of mucoadhesive drug delivery systems as targeting agents. European Journal of Pharmaceutics and Biopharmaceutics 71, 519–528 (2009).

41. Tsuji, S., et al. Human intelectin is a novel soluble lectin that recognizes galactofuranose in carbohydrate chains of bacterial cell wall. J Biol Chem 276, 23456–23463 (2001).

42. Castillo, P.A., et al. An Experimental Approach to Rigorously Assess Paneth Cell α-Defensin (Defa) mRNA Expression in C57BL/6 Mice. Scientific Reports 9, 13115 (2019).

43. Bevins, C.L. & Salzman, N.H. Paneth cells, antimicrobial peptides and maintenance of intestinal homeostasis. Nature Reviews Microbiology 9, 356–368 (2011).

44. Cochran, K.E., Lamson, N.G. & Whitehead, K.A. Expanding the utility of the dextran sulfate sodium (DSS) mouse model to induce a clinically relevant loss of intestinal barrier function. PeerJ 8, e8681–e8681 (2020).

45. Bashir, M.E.H., Louie, S., Shi, H.N. & Nagler-Anderson, C. Toll-Like Receptor 4 Signaling by Intestinal Microbes Influences Susceptibility to Food Allergy. The Journal of Immunology 172, 6978–6987 (2004).

46. Sorobetea, D., Holm, J.B., Henningsson, H., Kristiansen, K. & Svensson-Frej, M. Acute infection with the intestinal parasite Trichuris muris has long-term consequences on mucosal mast cell homeostasis and epithelial integrity. Eur J Immunol 47, 257–268 (2017).

47. Bramhall, M. & Zaph, C. Mastering gut permeability: New roles for old friends. Eur J Immunol 47, 236–239 (2017).

48. Atarashi, K., et al. Induction of colonic regulatory T cells by indigenous Clostridium species. Science 331, 337–341 (2011).

49. Lopetuso, L.R., Scaldaferri, F., Petito, V. & Gasbarrini, A. Commensal Clostridia: leading players in the maintenance of gut homeostasis. Gut Pathogens 5, 23 (2013).

50. Byndloss, M.X., et al. Microbiota-activated PPAR-γ signaling inhibits dysbiotic Enterobacteriaceae expansion. Science 357, 570–575 (2017).

51. Ostanin, D.V., et al. T cell transfer model of chronic colitis: concepts, considerations, and tricks of the trade. Am J Physiol Gastrointest Liver Physiol 296, G135–146 (2009).

52. Gupta, R.S., et al. The Public Health Impact of Parent-Reported Childhood Food Allergies in the United States. Pediatrics 142, e20181235 (2018).

53. Gupta, R.S., et al. Prevalence and Severity of Food Allergies Among US Adults. JAMA Netw Open 2, e185630–e185630 (2019).

54. Folkerts, J., et al. Butyrate inhibits human mast cell activation via epigenetic regulation of FcεRI-mediated signaling. Allergy 75, 1966–1978 (2020).

55. Nair, A.B. & Jacob, S. A simple practice guide for dose conversion between animals and human. J Basic Clin Pharm 7, 27–31 (2016).

56. Vernero, M., et al. The Usefulness of Microencapsulated Sodium Butyrate Add-On Therapy in Maintaining Remission in Patients with Ulcerative Colitis: A Prospective Observational Study. Journal of clinical medicine 9, 3941 (2020).

57. Facchin, S., et al. Microbiota changes induced by microencapsulated sodium butyrate in patients with inflammatory bowel disease. Neurogastroenterology & Motility 32, e13914 (2020).

58. Cai, R., Li, R., Qian, J., Xie, A. & Nie, K. The morphology and fabrication of nanostructured micelle by a novel block copolymer with linear–dendritic structure. Materials Science and Engineering: C 33, 2070–2077 (2013).

59. Megías, C., et al. Stability of sunflower protein hydrolysates in simulated gastric and intestinal fluids and Caco-2 cell extracts. LWT - Food Science and Technology 42, 1496–1500 (2009).

60. Fu, T.J., Abbott, U.R. & Hatzos, C. Digestibility of food allergens and nonallergenic proteins in simulated gastric fluid and simulated intestinal fluid-a comparative study. J Agric Food Chem 50, 7154–7160 (2002).

61. Law, C.W., Chen, Y., Shi, W. & Smyth, G.K. voom: precision weights unlock linear model analysis tools for RNA-seq read counts. Genome Biology 15, R29 (2014).

62. Wehkamp, J., et al. Reduced Paneth cell alpha-defensins in ileal Crohn’s disease. Proc Natl Acad Sci U S A 102, 18129–18134 (2005).

63. Nonnecke, E.B., et al. Human intelectin-1 (ITLN1) genetic variation and intestinal expression. Sci Rep 11, 12889 (2021).

64. Wehkamp, J., et al. Paneth cell antimicrobial peptides: topographical distribution and quantification in human gastrointestinal tissues. FEBS letters 580, 5344–5350 (2006).

65. Almalki, F., et al. Extensive variation in the intelectin gene family in laboratory and wild mouse strains. Sci Rep 11, 15548 (2021).

66. Callahan, B.J., et al. DADA2: High-resolution sample inference from Illumina amplicon data. Nature Methods 13, 581–583 (2016).

67. Wang, Q., Garrity George, M., Tiedje James, M. & Cole James, R. Naïve Bayesian Classifier for Rapid Assignment of rRNA Sequences into the New Bacterial Taxonomy. Applied and Environmental Microbiology 73, 5261–5267 (2007).

68. McMurdie, P.J. & Holmes, S. phyloseq: An R Package for Reproducible Interactive Analysis and Graphics of Microbiome Census Data. PLOS ONE 8, e61217 (2013).

69. Segata, N., et al. Metagenomic biomarker discovery and explanation. Genome Biology 12, R60 (2011).

70. Yang, C. Microbiome R package: microbiome biomarker analysis toolkit. R package version 0.99.0. (2020).

71. Turner, S., Pryer, K.M., Miao, V.P.W. & Palmer, J.D. Investigating Deep Phylogenetic Relationships among Cyanobacteria and Plastids by Small Subunit rRNA Sequence Analysis1. Journal of Eukaryotic Microbiology 46, 327–338 (1999).

72. Amann, R.I., Ludwig, W. & Schleifer, K.H. Phylogenetic identification and in situ detection of individual microbial cells without cultivation. Microbiological Reviews 59, 143–169 (1995).

73. Matsuki, T., et al. Development of 16S rRNA-gene-targeted group-specific primers for the detection and identification of predominant bacteria in human feces. Applied and environmental microbiology 68, 5445–5451 (2002).

