## Supplementary information for "Microbial metabolite butyrate-prodrug polymeric micelles demonstrate therapeutic efficacy in pre-clinical models of food allergy and colitis"

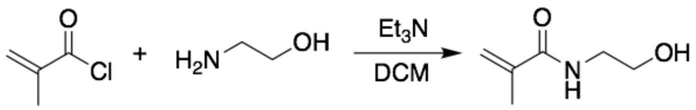


1 2

**Scheme S1** Synthesis of N-(2-hydroxyethyl) methacrylamide (HEMA, **2**)

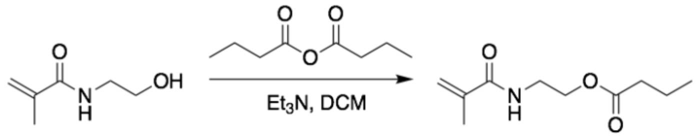


2 3

**Scheme S2** Synthesis of N-(2-butanoyloxyethyl) methacrylamide (BMA, **3**)

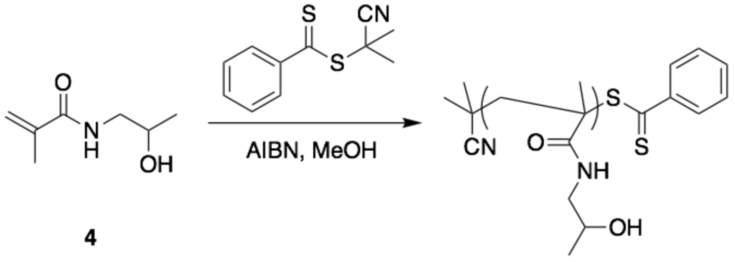


4

**5 (pHPMA)**

**Scheme S3** Synthesis of poly(2-hydroxypropyl methacrylamide) (pHPMA, **5**)

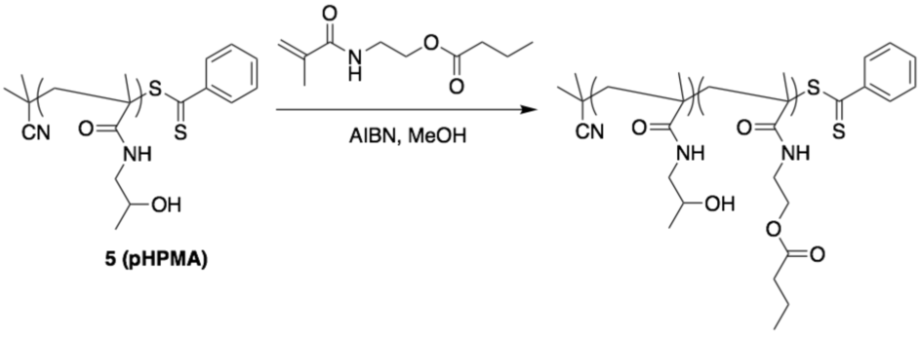


**5 (pHPMA)**

**6 (pHPMA-b-pBMA)**

**Scheme S4** Synthesis of pHPMA-*b*-pBMA (**6**)

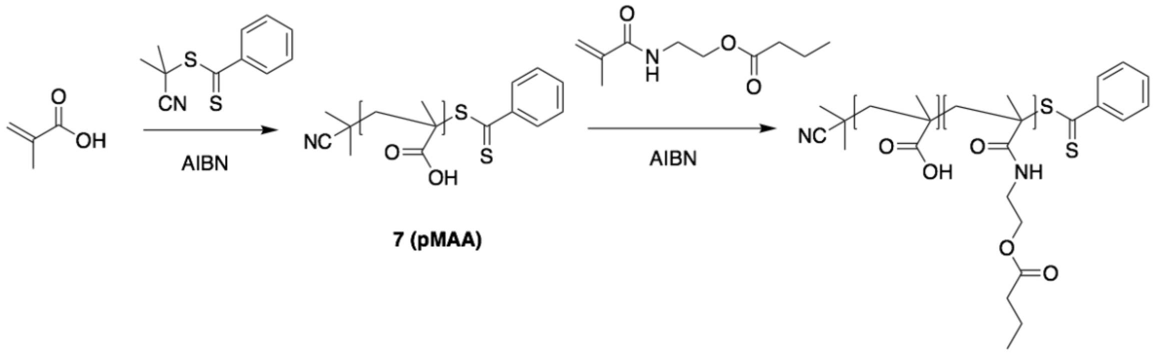


**7 (pMAA)**

**8 (pMAA-*b*-pBMA)**

**Scheme S5** Synthetic route of pMAA-*b*-pBMA (**8**)

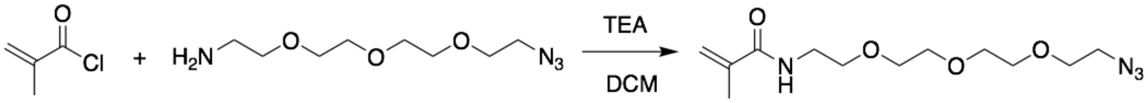


**9** $N_3$-PEG$_4$-MA

**Scheme S6** Synthesis of hydrophilic azide monomer $N_3$-PEG$_4$-MA (**9**)

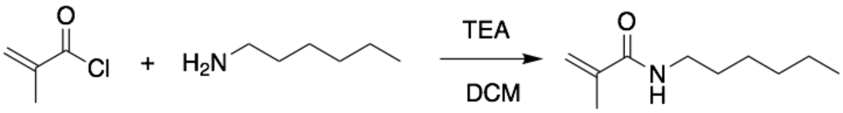


**10**

**Scheme S7** Synthesis of hydrophobic control monomer N-hexyl methacrylamide (**10**)

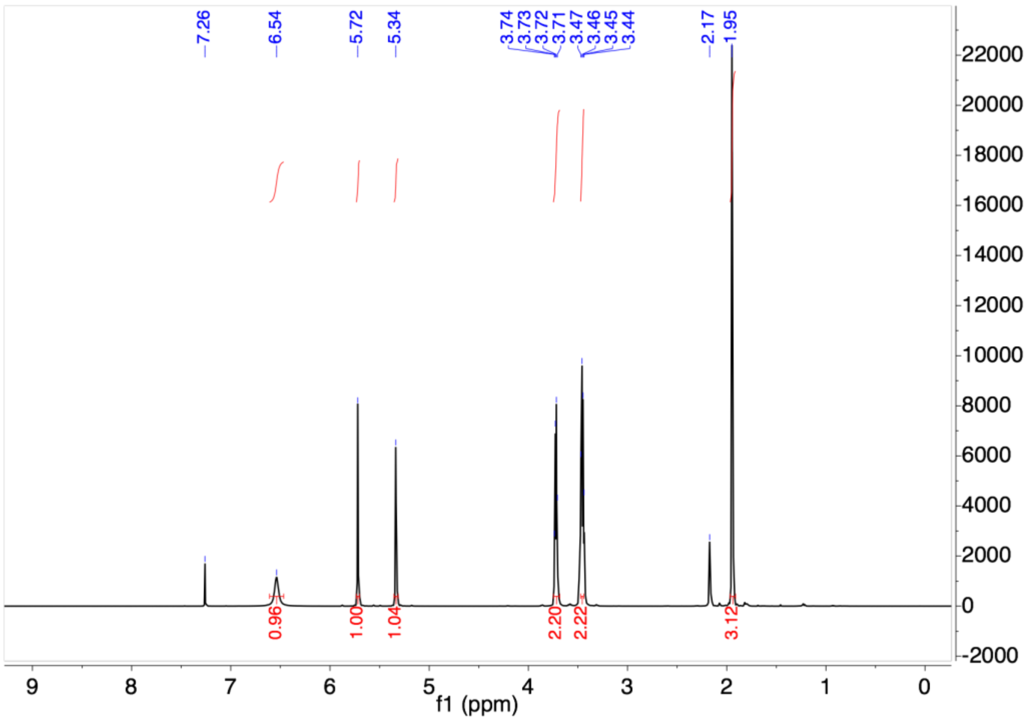

**Fig. S1.** $^1$H-NMR (500 MHz, $CDCl_3$) of HEMA (**2**)

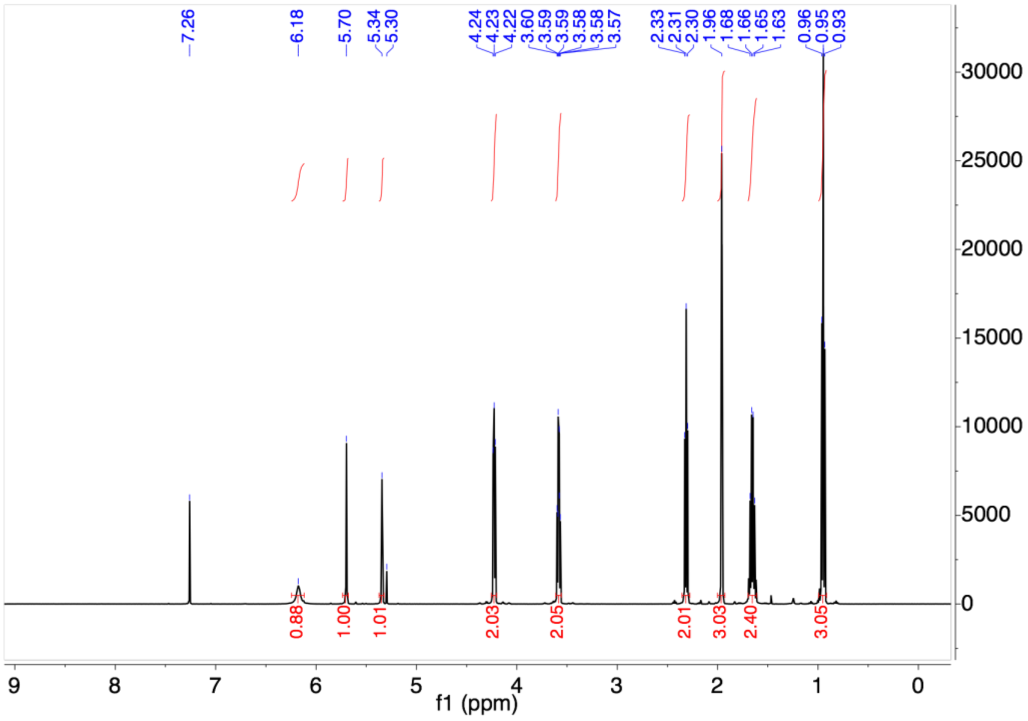

**Fig. S2.** $^1$H-NMR (500 MHz, $CDCl_3$) of BMA (**3**)

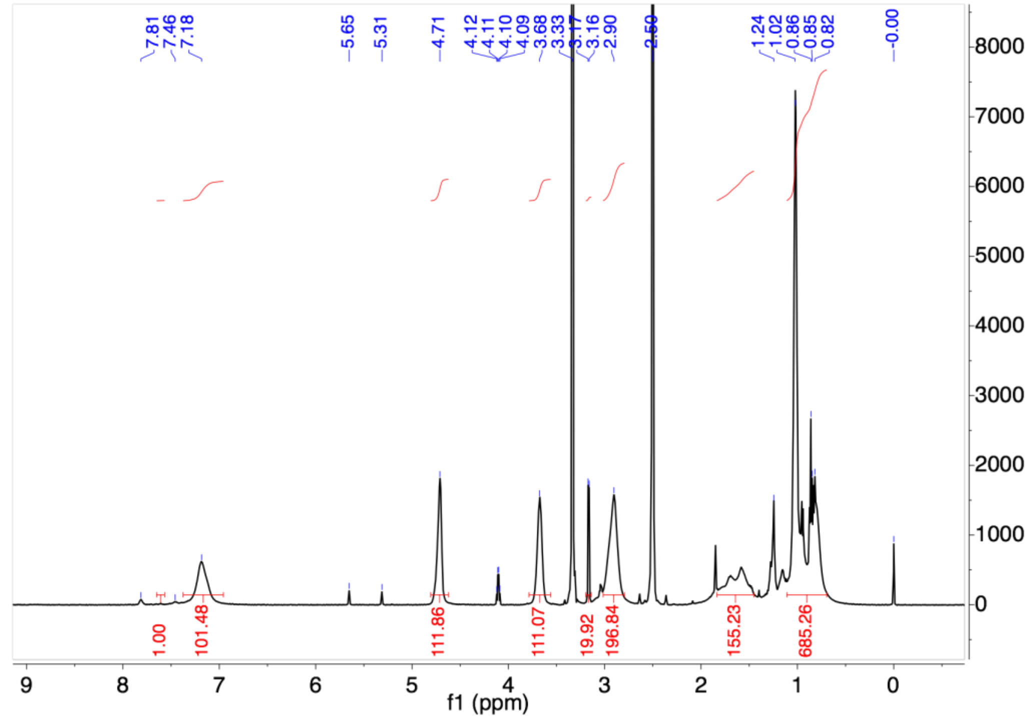

**Fig. S3.** $^{1}$H-NMR (500 MHz, DMSO-d6) of pHPMA (**5**)

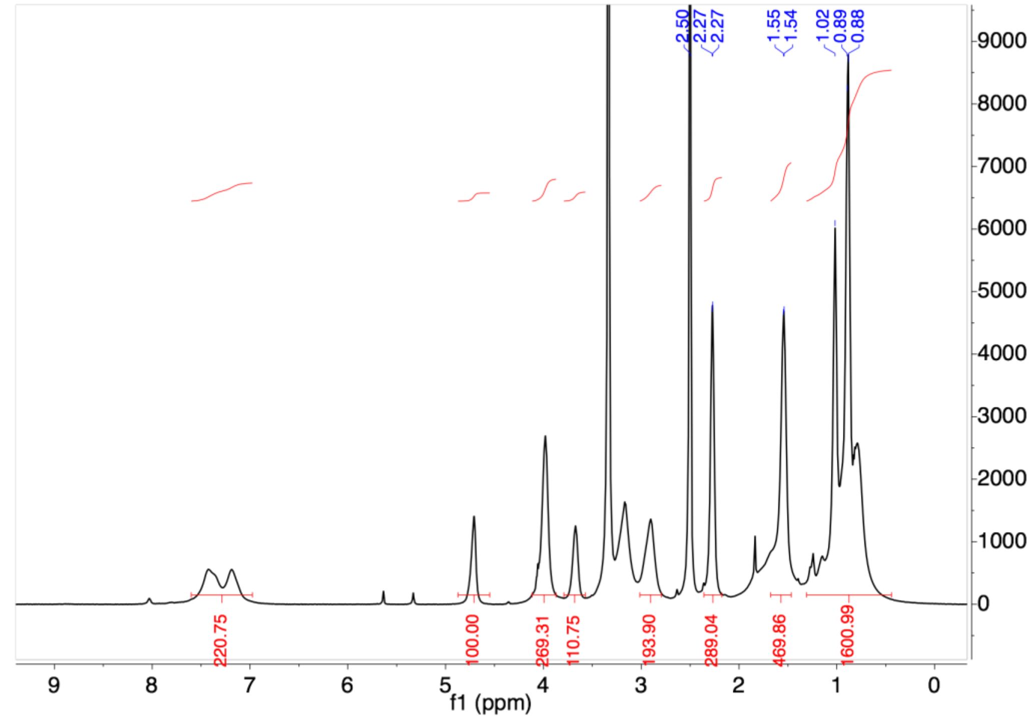

**Fig. S4.** $^{1}$H-NMR (500 MHz, DMSO-d6) of pHPMA-*b*-pBMA (**6**)

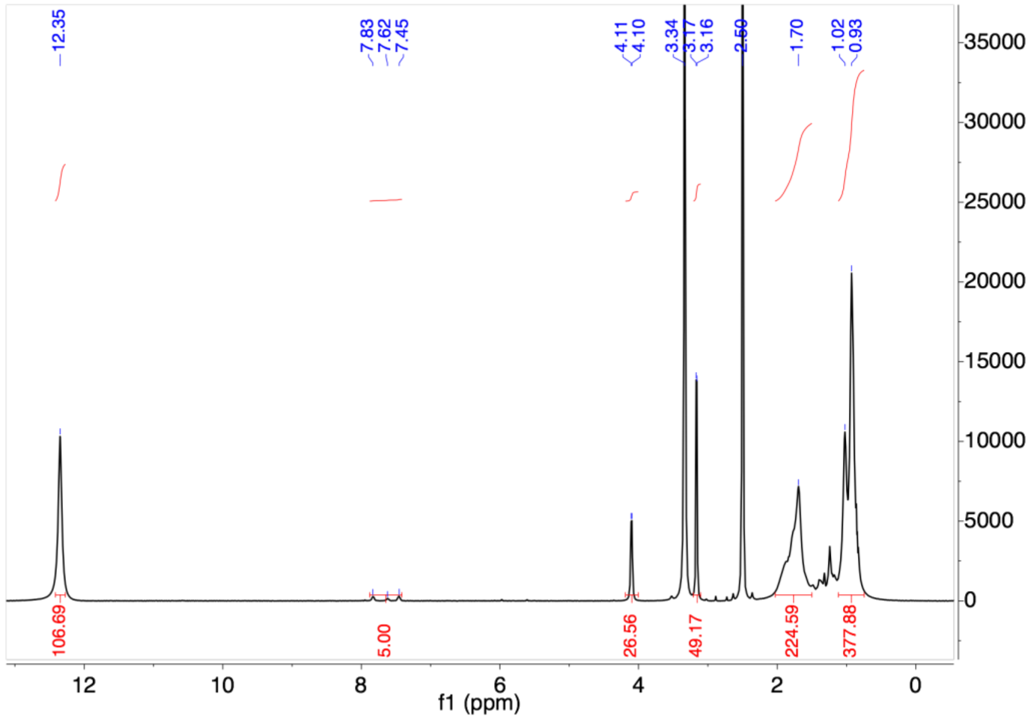


**Fig. S5.** $^{1}$H-NMR (500 MHz, DMSO-d6) of pMAA (**7**)

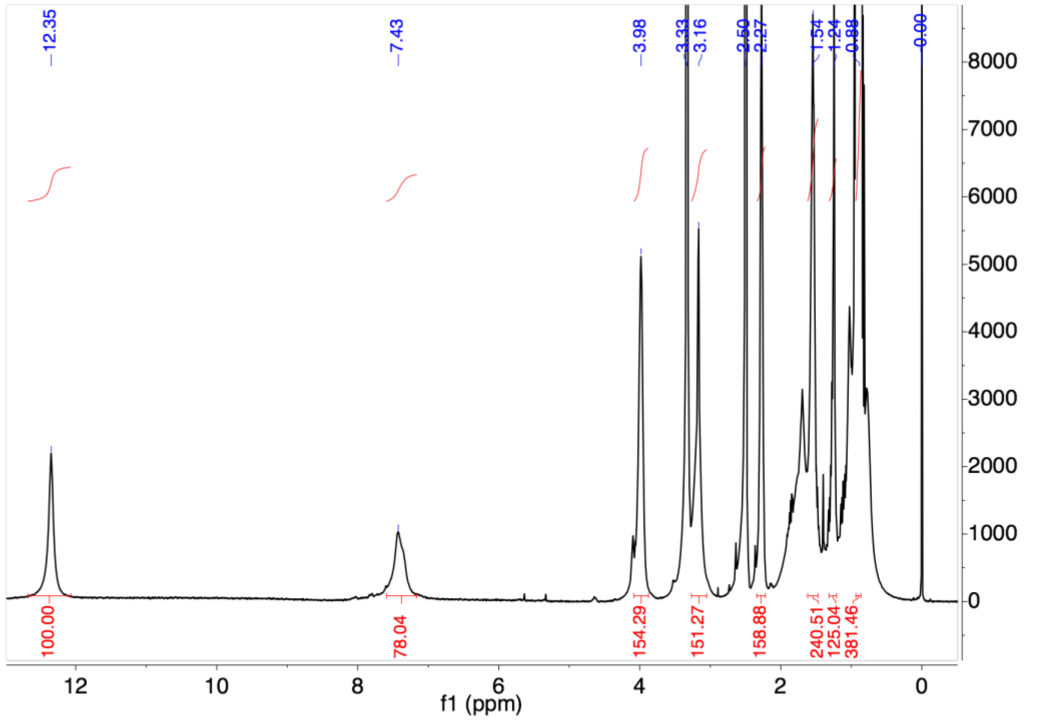


**Fig. S6.** $^{1}$H-NMR (500 MHz, DMSO-d6) of pMAA-*b*-pBMA (**8**)

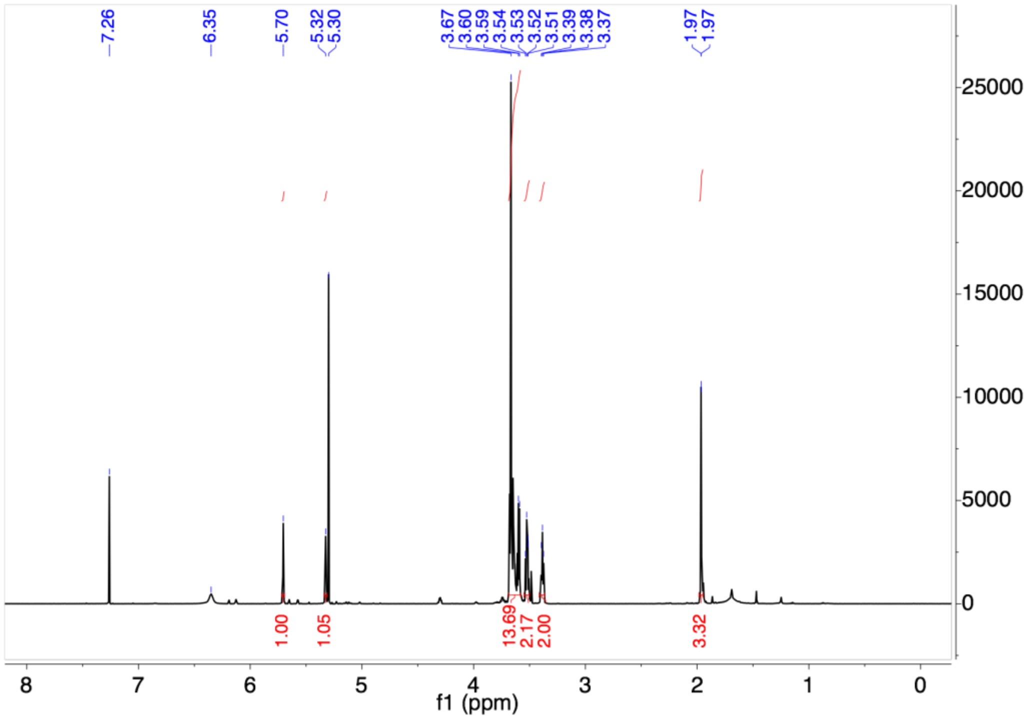


**Fig. S7.** $^{1}$H-NMR (500 MHz, $CDCl_3$) of $N_3$-$PEG_4$-MA (**9**)

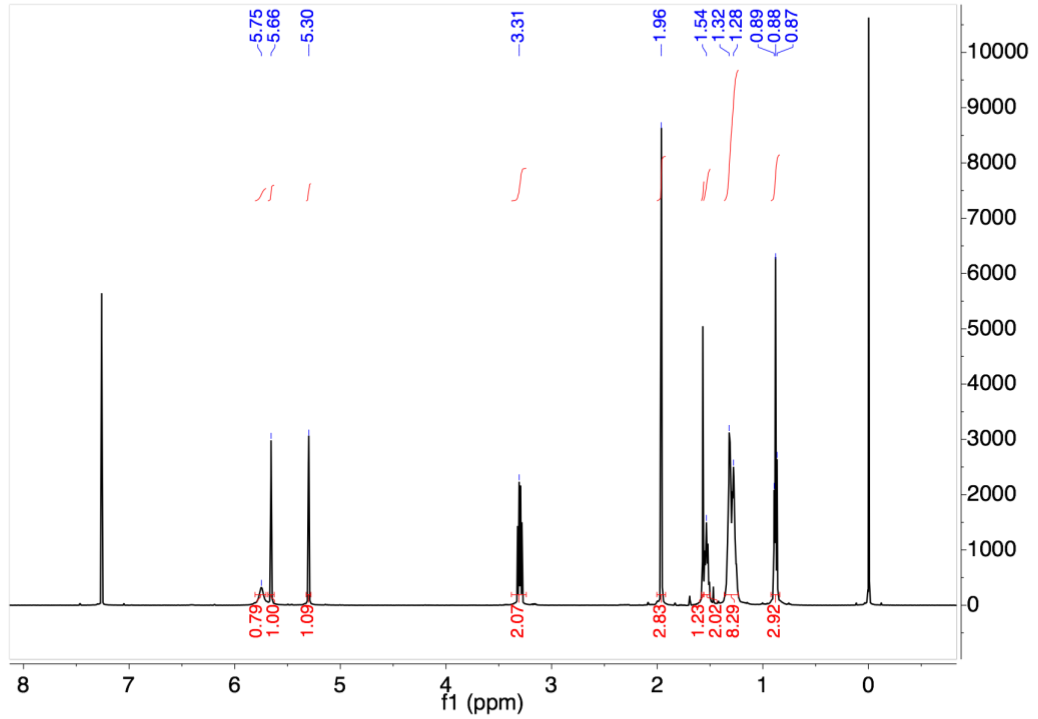


**Fig. S8.** $^{1}$H-NMR (500 MHz, $CDCl_3$) of N-hexyl methacrylamide (**10**)

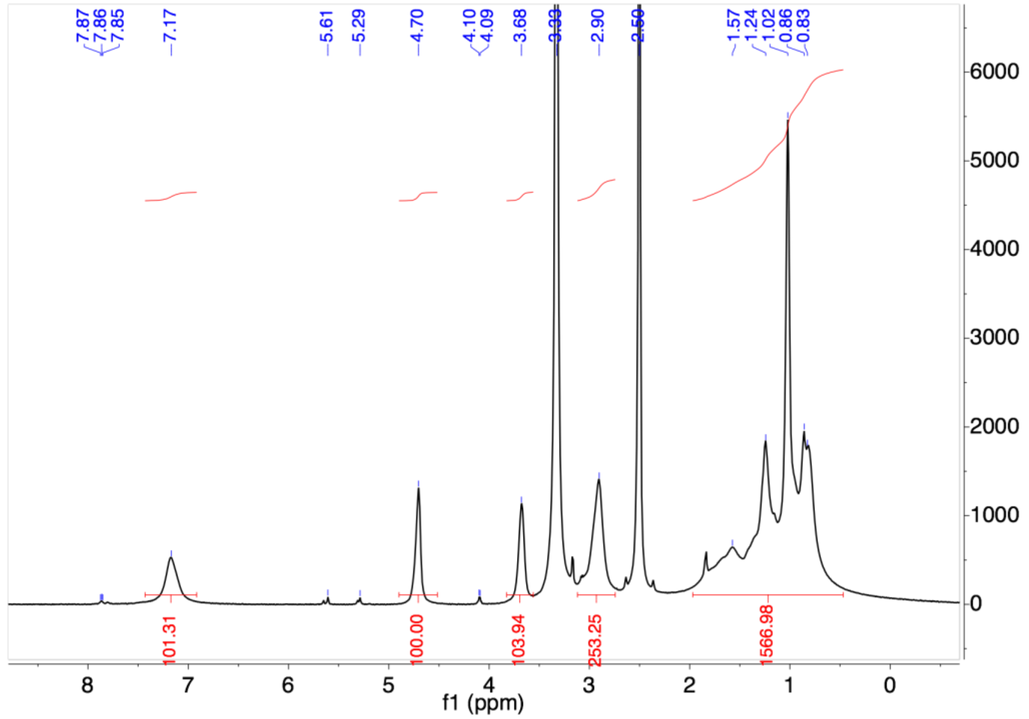


**Fig. S9.** $^{1}$H-NMR (500 MHz, DMSO-d6) of control polymer pHPMA-*b*-pHMA

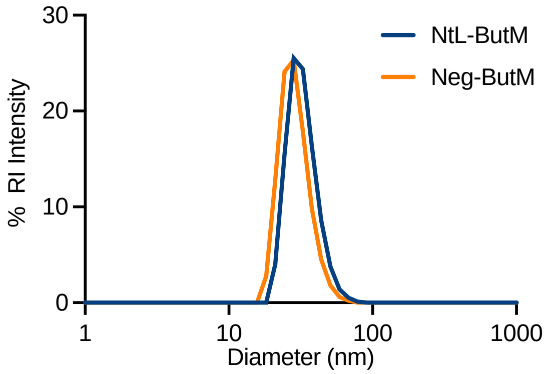


**Fig. S10.** Dynamic light scattering (DLS) shows that the micelles NtL-ButM and Neg-ButM have similar hydrodynamic diameters, at sub-hundred nanometer.

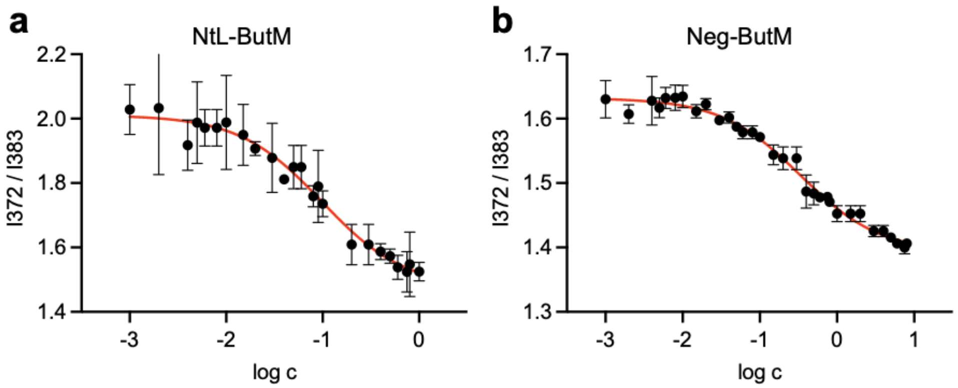


**Fig. S11.** Critical micelle concentrations (CMC) of NtL-ButM (**a**) and Neg-ButM (**b**) measured by pyrene fluorescent intensity of peak 1 over peak 3. The CMC was determined by the IC50 fitted by a sigmoidal curve.

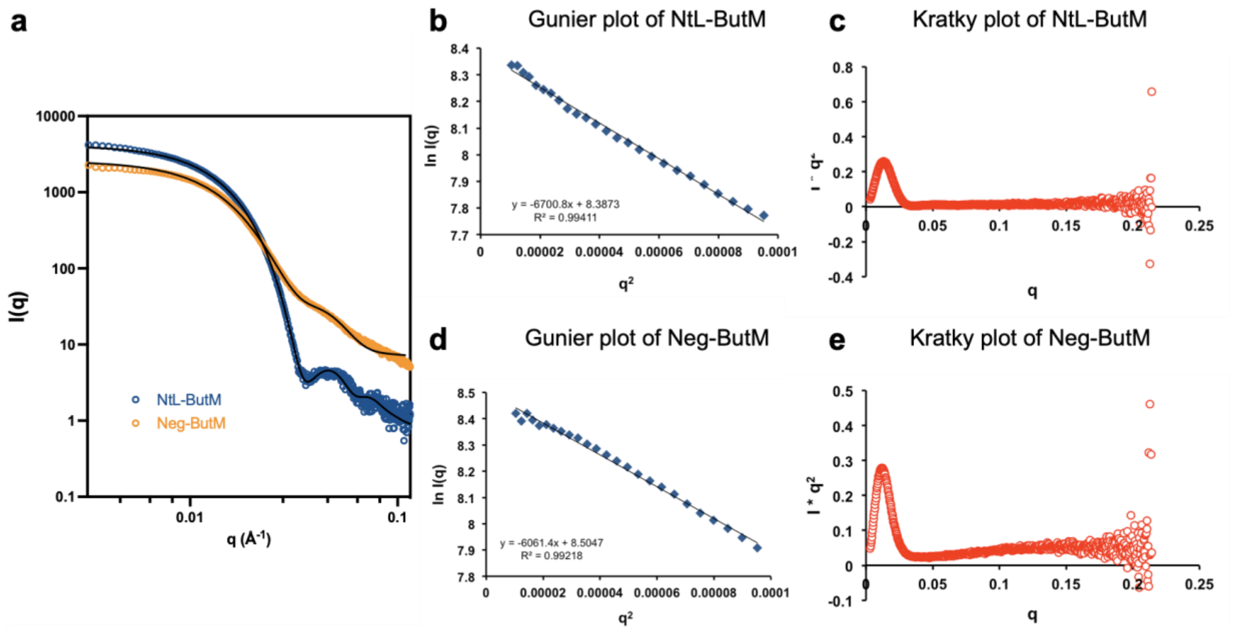


**f**

| | *c* (mg mL$^{-1}$) | radius$_{core}$ (Å) | thickness$_{shell}$ (Å) | SLD$_{core}$ ($10^{-6}$ Å$^{-2}$) | SLD$_{shell}$ ($10^{-6}$ Å$^{-2}$) | volume fraction |
|---|---|---|---|---|---|---|
| NtL-ButM | 2.0 | 119.31 ± 0.16 | 72.63 ± 0.31 | 16.1821 ± 0.0016 | 10.5338 ± 0.0003 | 0.007381 |
| Neg-ButM | 2.0 | 83.84 ± 0.34 | 98.97 ± 0.22 | 16.9295 ± 0.0023 | 10.8700 ± 0.0004 | 0.01393 |

**g**

| | *c* (mg mL$^{-1}$) | *d* (nm) | *N* (×$10^{14}$ mL$^{-1}$) | *Mw* (×$10^{6}$ g mol$^{-1}$) | $N_{agg}$ |
|---|---|---|---|---|---|
| NtL-ButM | 2.0 | 158.9 | 2.492 | 4.831 | **119** |
| Neg-ButM | 2.0 | 122.5 | 5.441 | 2.213 | **92** |

**Fig. S12.** Small-angle X-ray scattering (SAXS) characterization of NtL-ButM and Neg-ButM micelles. **a**, SAXS data of NtL-ButM and Neg-ButM. Data are fitted with polydisperse core-shell model. **b**, Gunier plot (ln($q$) vs. $q^2$) of NtL-ButM revealed the radius of gyration of the micelle. **c,** Kratky plot ($I\ q^2$ vs. $q$) of NtL-ButM revealed the spherical structure if the micelle. **d,** Gunier plot of Neg-ButM micelle. **e**, Kratky plot of Neg-ButM micelle. **f**, Table of fitting parameters of NtL-ButM and Neg-ButM using a polydisperse core-shell sphere model. **g,** Table of the mean distance between micelles *d*, number of micelles per unit volume *N*, molecular weight of the micelle *Mw*, and the aggregation number $N_{agg}$, calculated from the fitting parameters of a polydisperse core-shell sphere model.

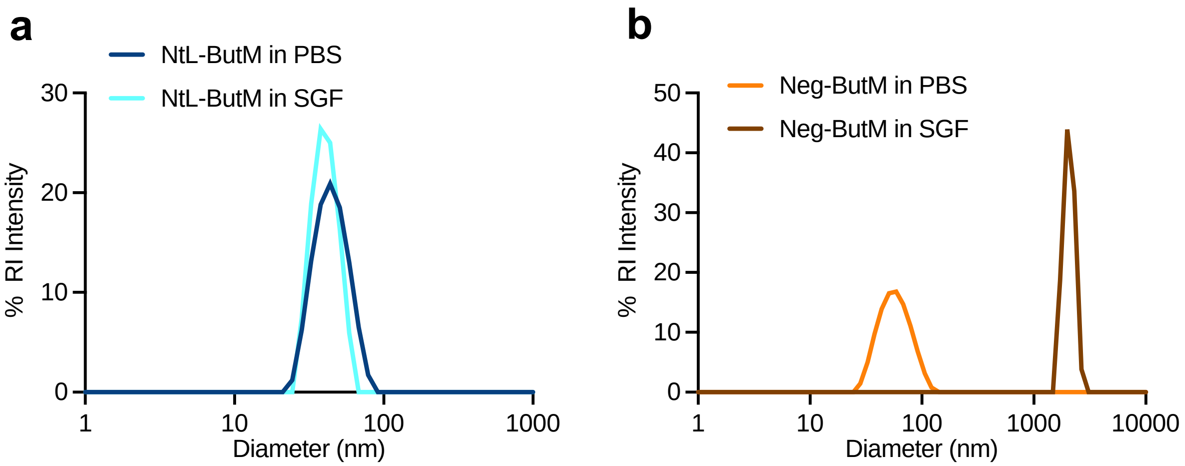


**Fig. S13.** The DLS data of diameter distribution (by intensity) of micelles NtL-ButM (**a**) and Neg-ButM (**b**) in PBS or simulated gastric fluid (SGF).

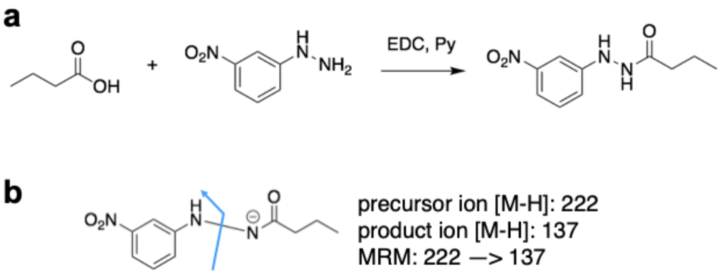


**Fig. S14.** Derivatization of butyrate for LC-MS/MS analysis and the release of butyrate from NtL-ButM/Neg-ButM in simulated gastric/intestinal fluids. **a,** Derivatization reaction of butyrate with 3-nitrophenylhydrazine (NPH) to generate UV active butyrate-NPH. **b,** The multiple reaction monitoring (MRM) of 222 → 137 was used to quantify butyrate-NPH in LC-MS/MS.

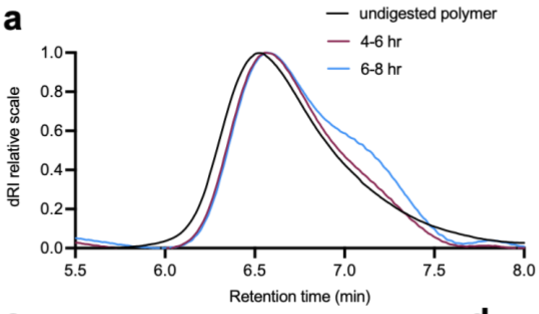


b

| | Fecal sample collected from 4-6 hr | Fecal sample collected from 6-8 hr |
|---|---|---|
| Mn | 37.17 kDa | 34.21 kDa |
| Mw | 47.24 kDa | 44.63 kDa |
| PDI | 1.271 | 1.305 |
| Mn loss | -20.03% | -26.40% |

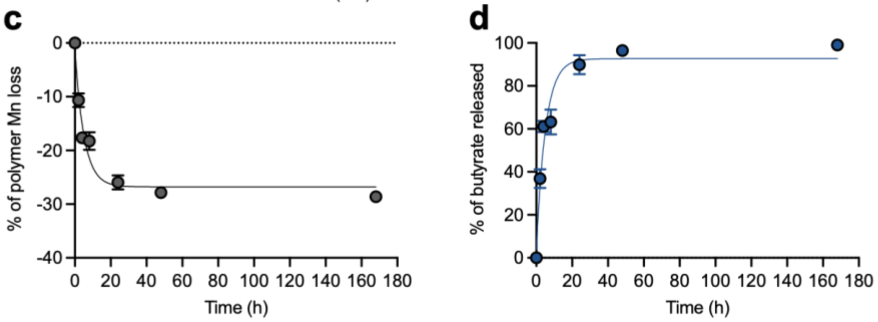


**Fig. S15.** Stability of pHPMA-b-pBMA polymer *in vitro* or *in vivo*. **a**, Gel permeation chromatography (GPC) elution profiles (measured by differential refractive index (dRI) over time) of polymers collected from pooled fecal samples of two mice treated with NtL-ButM at 4-6 hr (red) or 6-8 hr (blue) post-gavage. Black curve: polymer control. **b,** The table of molecular weight of digested polymer measured by GPC, including the number averaged molecular weight (Mn) and weight averaged molecular weight (Mw), with their polydispersity index (PDI) and the Mn loss compared to undigested polymer control. **c, d,** The Mn loss of pHPMA-b-pBMA polymer in 125 mM NaOH solution over 7 days (**c**), or the percentage of butyrate released from the polymer (**d**), measured by GPC, n=3, Data represent mean ± s.e.m.

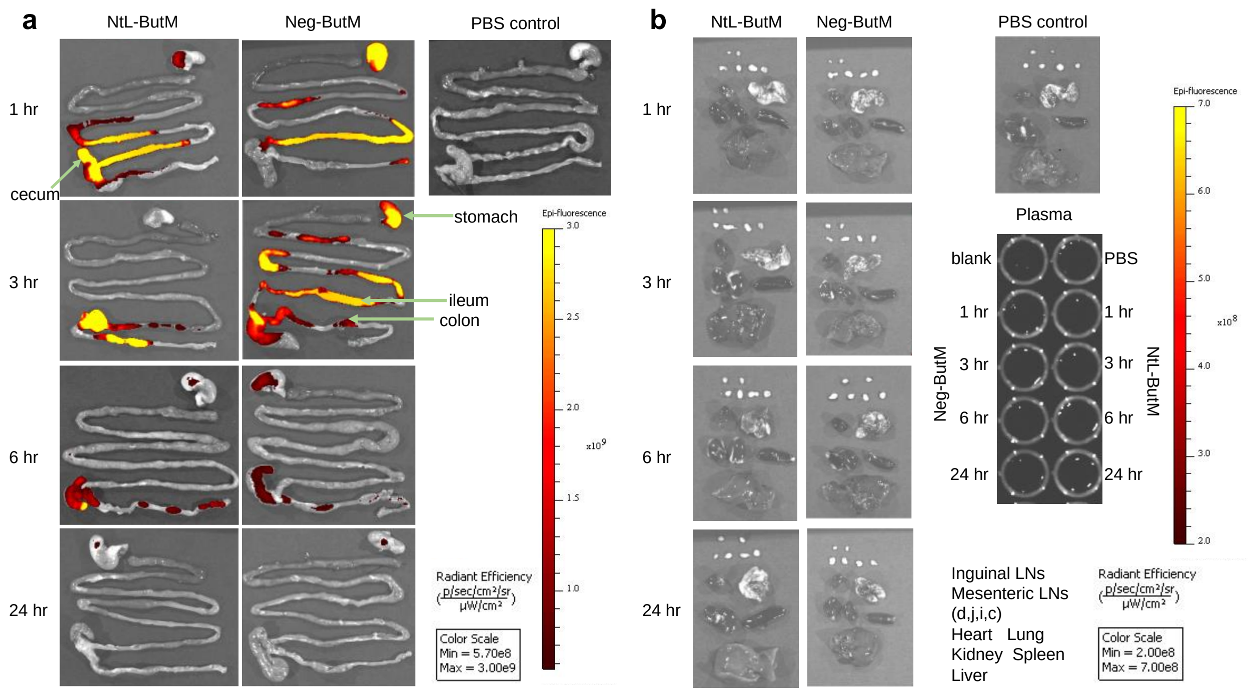


**Fig. S16.** The biodistribution of NtL-ButM or Neg-ButM in the gastrointestinal (GI) tract of non-antibiotic treated SPF mice. Mice were administered IR750 labeled NtL-ButM or Neg-ButM intragastrically and euthanized at the indicated time points to measure the fluorescent signals in the GI tract (**a**), and other major organs and plasma (**b**) using an in vivo imaging system (IVIS). Both polymers were chemically modified with azide and labeled with dye IR750. After a single oral administration of labeled NtL-ButM or Neg-ButM (one mouse per time point per treatment group), IVIS showed Neg-ButM retained in the stomach for more than 6 hr while NtL-ButM moved to the cecum quickly. Both polymers were cleared from the GI tract after 24 hr, and there was no absorption of either butyrate micelle into the systemic circulation. Mesenteric LNs (d, duodenum-draining; j, jejunum-draining; I, ileum-draining; c, colon-draining).

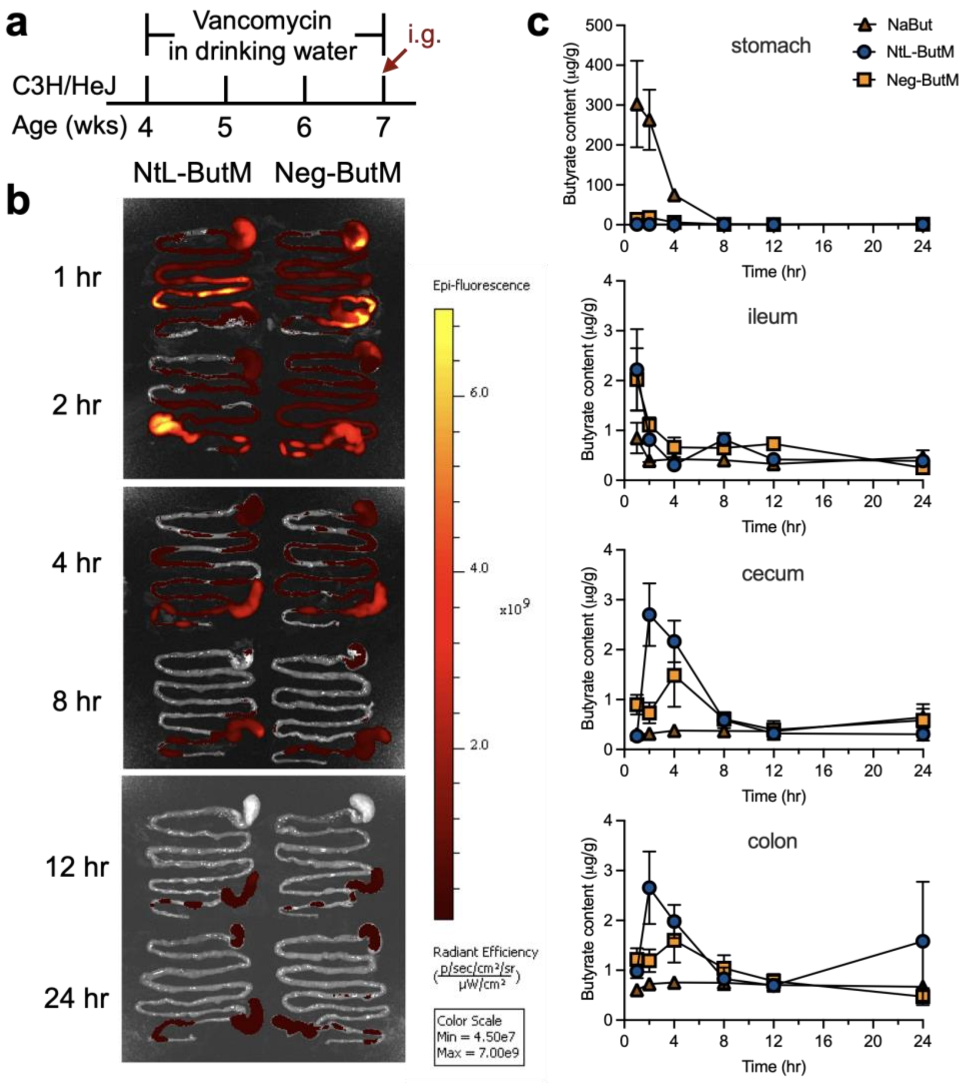


**Fig. S17.** The biodistribution and *in vivo* butyrate release of NtL-ButM or Neg-ButM in the GI tract of vancomycin-treated mice. **a,** Butyrate producing bacteria were depleted by treating 4 week old SPF C3H/HeJ mice with 200 mg/L vancomycin in the drinking water for three weeks prior to intragastric (i.g.) administration of each of the butyrate formulations. **b,** Mice were administered IR750 labeled NtL-ButM or Neg-ButM intragastrically and euthanized at the indicated time points to measure the fluorescent signals in the GI tract by IVIS. **c,** The amount of butyrate released in the stomach, ileum, cecum, or colon after a signal intragastric administration of sodium butyrate, NtL-ButM, or Neg-ButM (at an equivalent butyrate dose of 2.24 mg/g) to vancomycin-treated mice. At the indicated time points, the stomach, ileum, cecum, and colon were harvested, and the whole tissues were homogenized, followed by butyrate extraction and quantification by LC-MS/MS. n = 5 mice per group per time point. Data represent mean ± s.e.m.

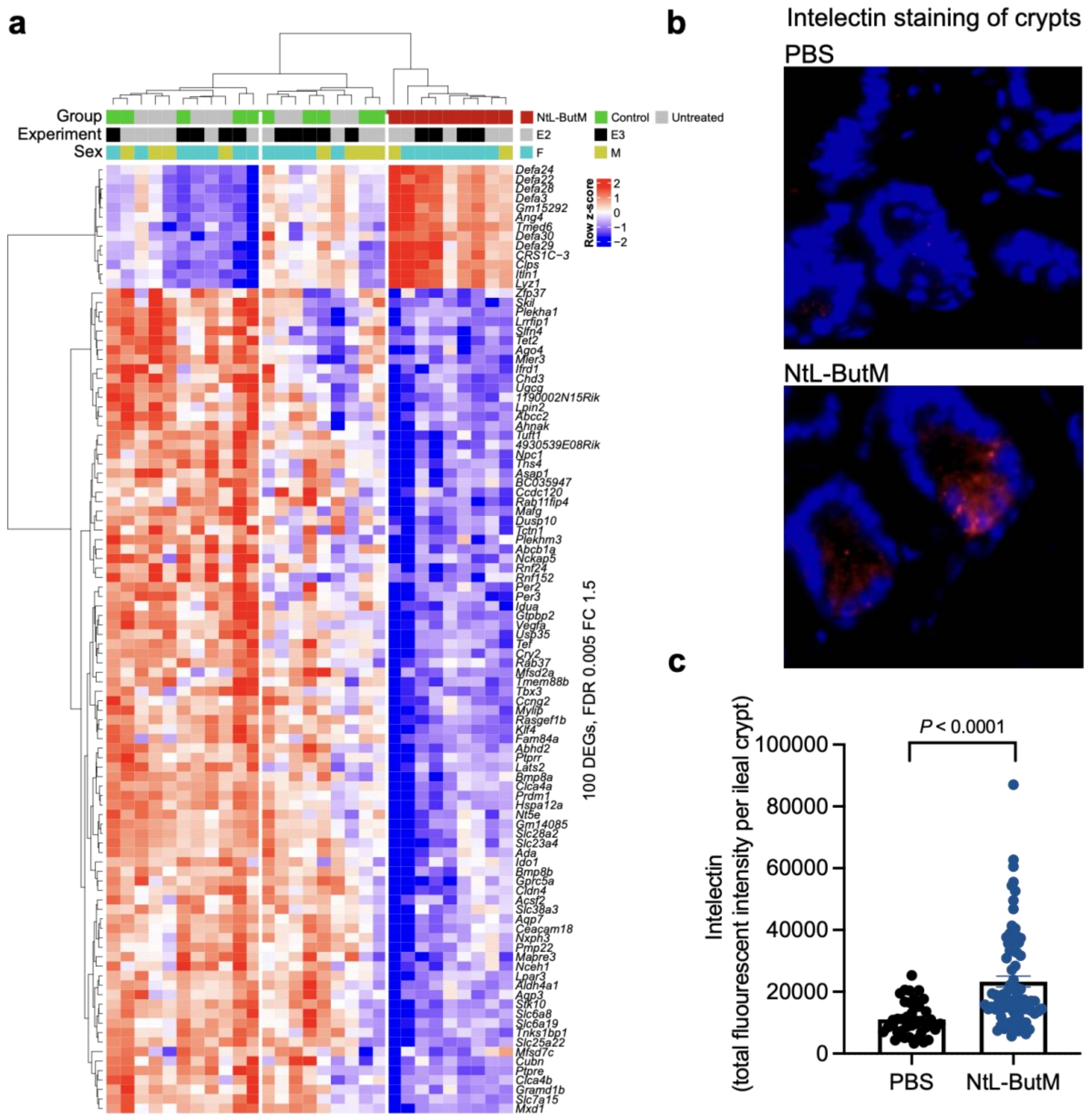


**Fig. S18.** NtL-ButM induced an ileal gene expression signature that is almost entirely anti-microbial peptides (AMPs). **a,** One week of daily dosing of 0.8 mg/g NtL-ButM to germ-free (GF) C3H/HeN mice induces a unique gene expression signature in the ileum compared to untreated and inactive polymer controls as measured by RNA sequencing of isolated intestinal epithelial cells. Top 100 significant differentially expressed genes (DEGs) at False Discovery Rate (FDR)-adjusted $P$<0.005 and fold change (FC) ≥ 1.5 or ≤−1.5 are shown. Annotation bars of the three groups, experiment batches (E2 and E3), and gender (female, male) are shown above the heatmap. **b,** Fluorescent imaging of intelectin protein in small intestine sections from control or treated mice. Blue (DAPI), red (intelectin). **c,** Intelectin protein is quantified by total fluorescence signal per crypt of small intestine. n = 3 PBS-treated and 4 NtL-ButM treated mice, with >15 crypts quantified per mouse. Data represent mean ± s.e.m. limma voom with precision weights was used in **a**. Two-sided Student’s t-test was used in **c**.

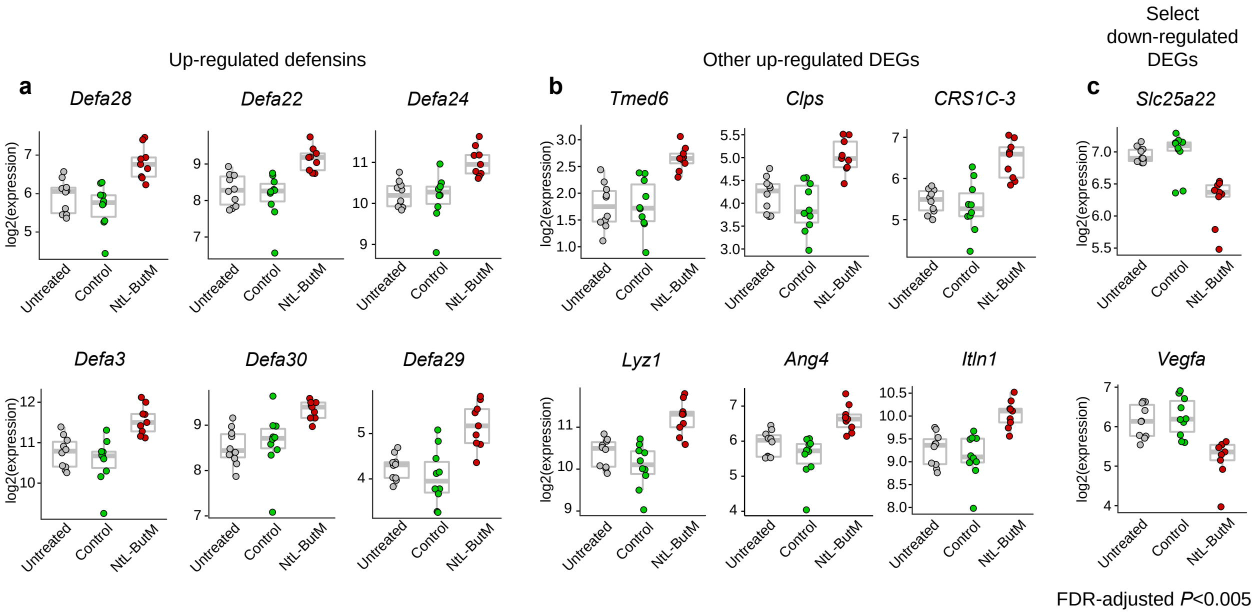


**Fig. S19.** Differentially expressed genes (DEGs) in the ileum of GF mice that were treated daily with 0.8 mg/g NtL-ButM for one week, compared to untreated and inactive polymer controls as measured by RNA sequencing of isolated ileal epithelial cells (see **Fig. S18**). The unit of the value is TMM-normalized and log2-transformed read counts.

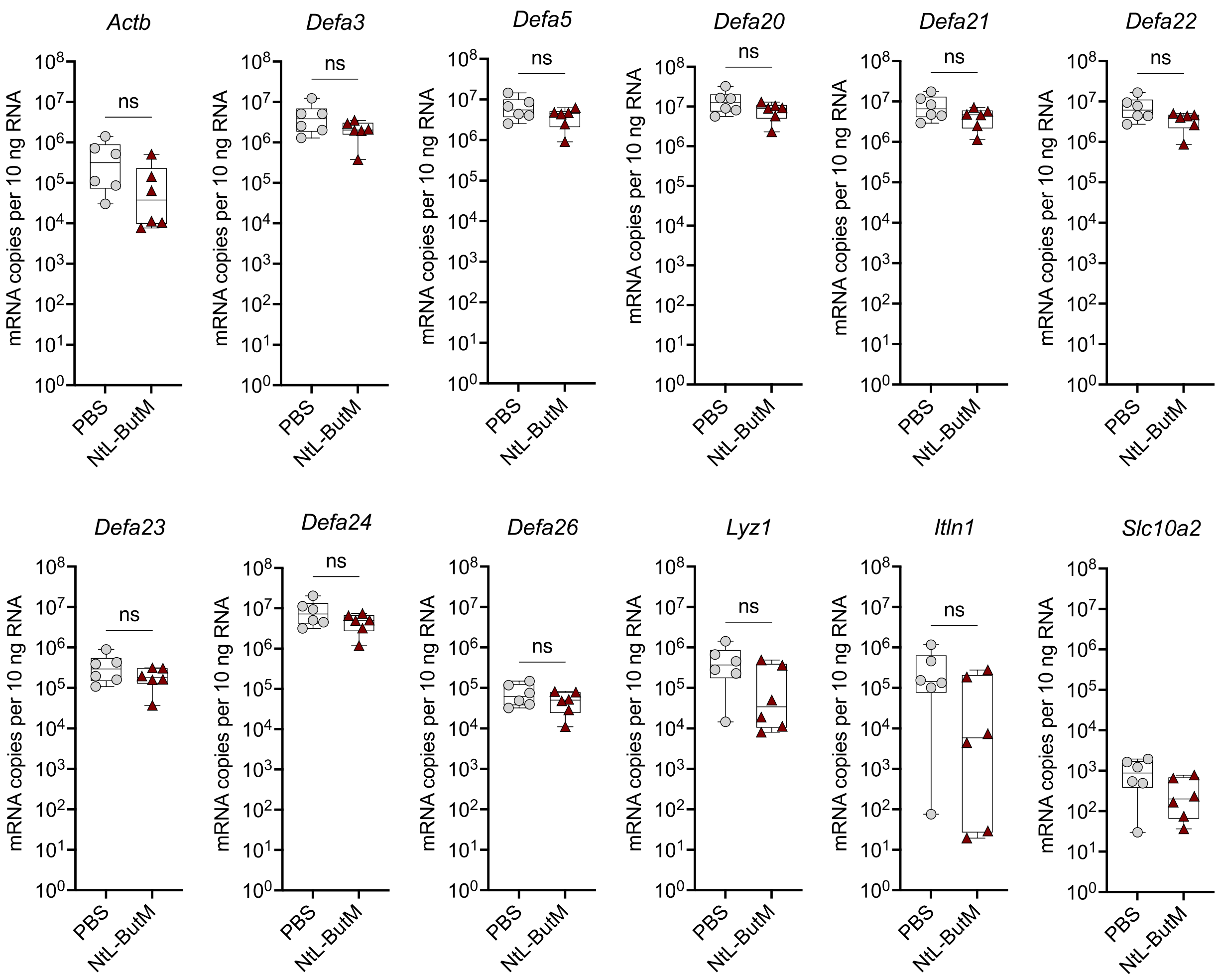


**Fig. S20.** NtL-ButM does not increase gene expression of defensins in SPF mice**.** Differentially expressed genes (DEGs) in the ileum of 8-week-old SPF C57BL/6 mice that were treated daily with 0.8 mg/g NtL-ButM (n = 6) for one week, compared to PBS-treated controls (n = 6) as measured by RT-qPCR of ileal epithelial cells. Data represent mean ± s.e.m. Data analyzed using two-sided Student's t-test.

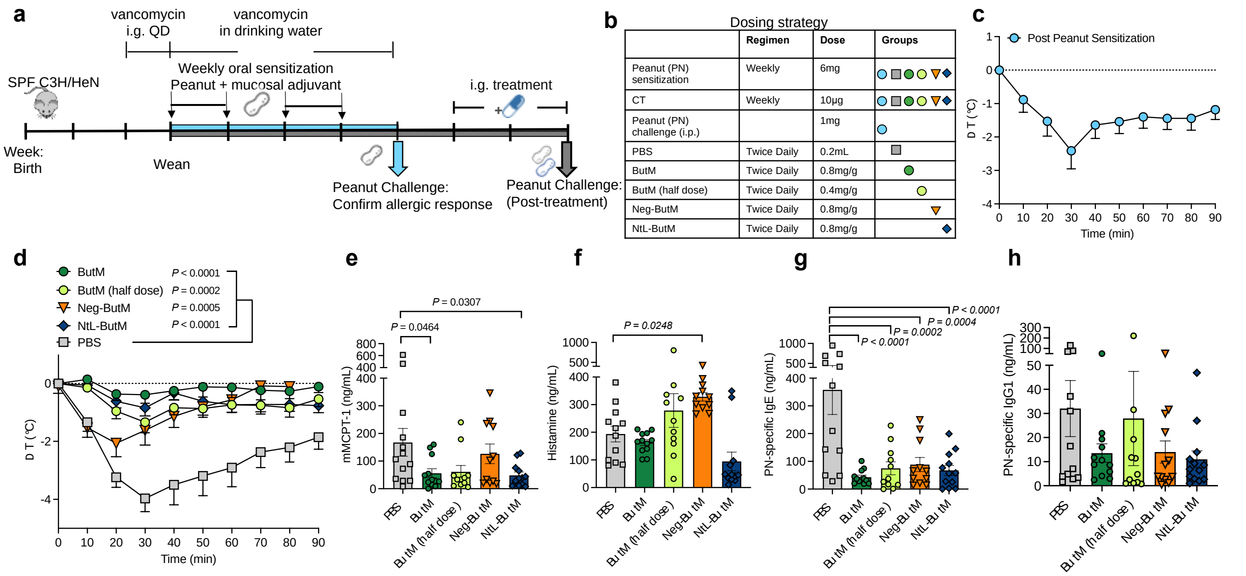


**Fig. S21.** Butyrate micelle treatment reduced the anaphylactic response to peanut challenge in a dose-dependent manner. **a, b,** Experimental schema, and the dosing strategy. All the mice were sensitized weekly by i.g. gavage of 6 mg of PN plus 10 μg of the mucosal adjuvant cholera toxin (CT). **c**, A uniform allergic response was confirmed by challenging one group of mice (n=7) with 1 mg of PN i.p. and measuring reduction in core body temperature as an indication of anaphylaxis. QD: once a day. **d-h**, The rest of mice were randomized into five treatment groups and received either PBS (n=12), ButM (n=12), ButM (half dose) (n=11), Neg-ButM only (n=11), or NtL-ButM only (n=12). **d,** Change in core body temperature following challenge with peanut extract. The area under curve (AUC) was compared among groups. **e-h**, Serum mMCPT-1 (**e**), histamine (**f**), peanut-specific IgE (**g**), and peanut-specific IgG1 (**h**) from mice in **d**. Data represent mean ± s.e.m. Data analyzed using one-way ANOVA with Dunnett's post-test.

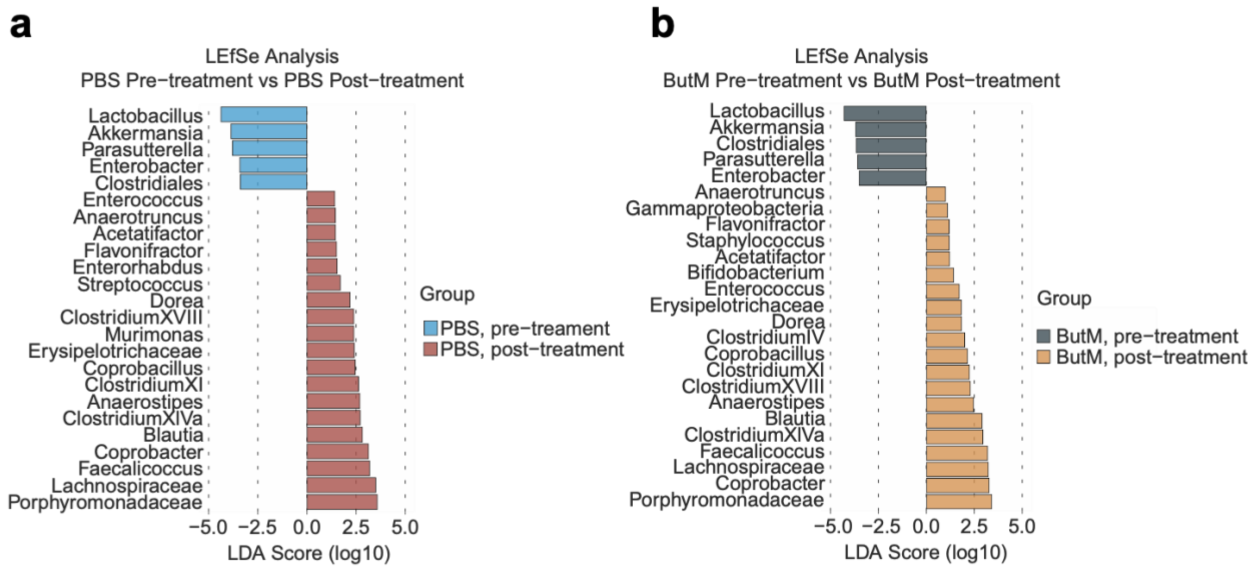


**Fig. S22.** Differentially abundant taxa within each treatment group before and after two-week treatment with PBS (**a**) or ButM (**b**) as analyzed by LEfSe from **Fig. 4**.

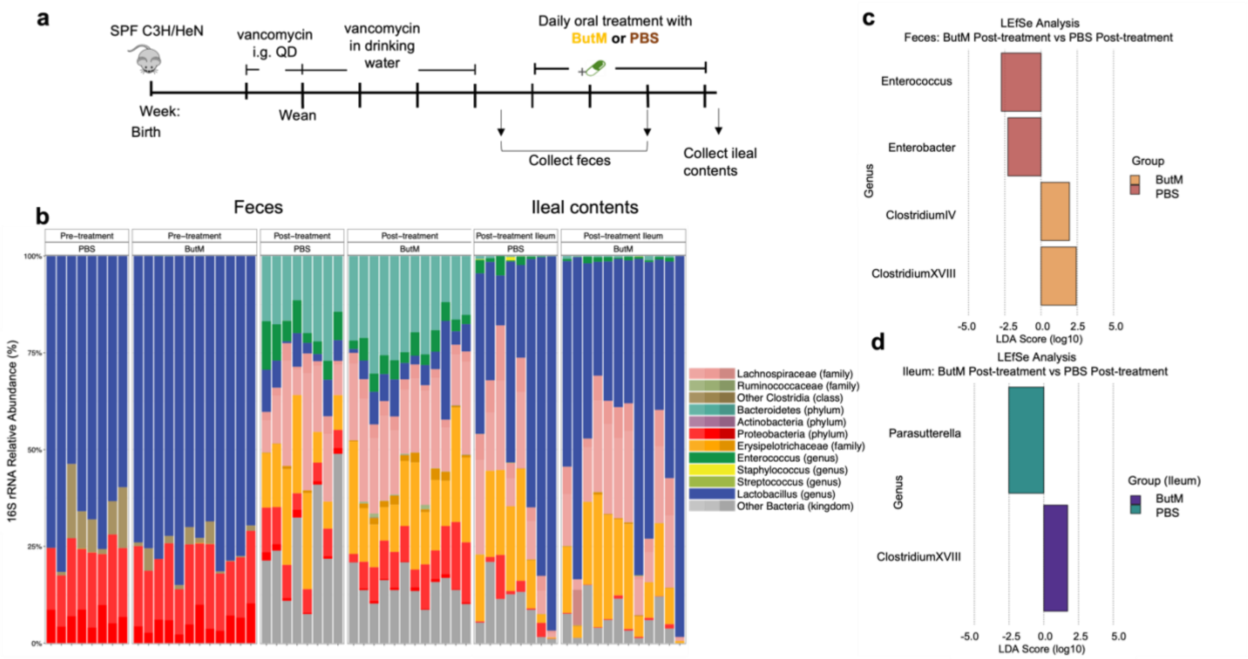


**Fig S23.** Treatment with ButM in unsensitized mice increases relative abundance of Clostridium clusters in the ileum and feces. **a**, Experimental schema. Mice are treated for five weeks with vancomycin, either by i.g. gavage pre-weaning or in the drinking water post-weaning. Upon cessation of vancomycin, mice were treated with PBS (*n* = 8) or 800mg/kg ButM (*n* = 12) twice daily. Ileal contents and fecal samples were collected just after ending antibiotic administration (pre-treatment) or after two weeks of treatment with PBS or ButM (post-treatment) at euthanasia (as in **Fig 4**). **b**, Relative abundance of bacterial taxa in feces and ileal contents. **c**, Differentially abundant taxa in feces between mice treated with PBS or ButM post-treatment analyzed by LEfSe. **d**, Differentially abundant taxa in ileal contents between mice treated with PBS or ButM post-treatment analyzed by LEfSe. QD: once a day.

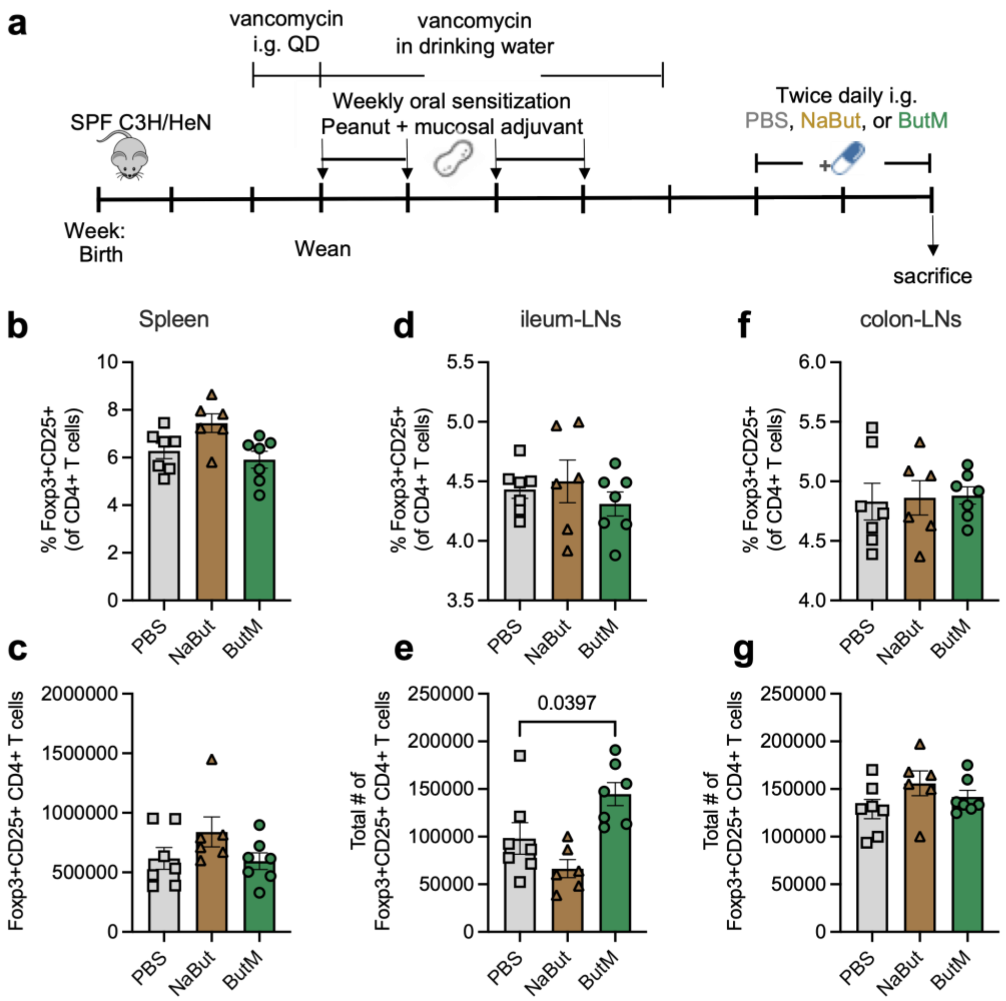


**Fig. S24.** Butyrate micelle treatment did not significantly affect regulatory T cell (Treg) populations in spleen or mesenteric LNs. **a,** Experimental schema. All of the vancomycin-treated mice were sensitized weekly by i.g. gavage of 6 mg of PN plus 10 μg of the mucosal adjuvant cholera toxin. The mice were then intragastrically treated with either PBS (n = 7), sodium butyrate (NaBut, n = 6), or ButM (n = 7) twice daily for two weeks. The mice were euthanized after treatment, and their spleen (**b,c**), ileum draining LNs (**d,e**), and colon draining LNs (**f,g**) were harvested and processed for flow cytometric analysis. Gating strategy is shown in **Fig. S25**. Data represent mean ± s.e.m. Data analyzed using one-way ANOVA with Dunnett's post-test.

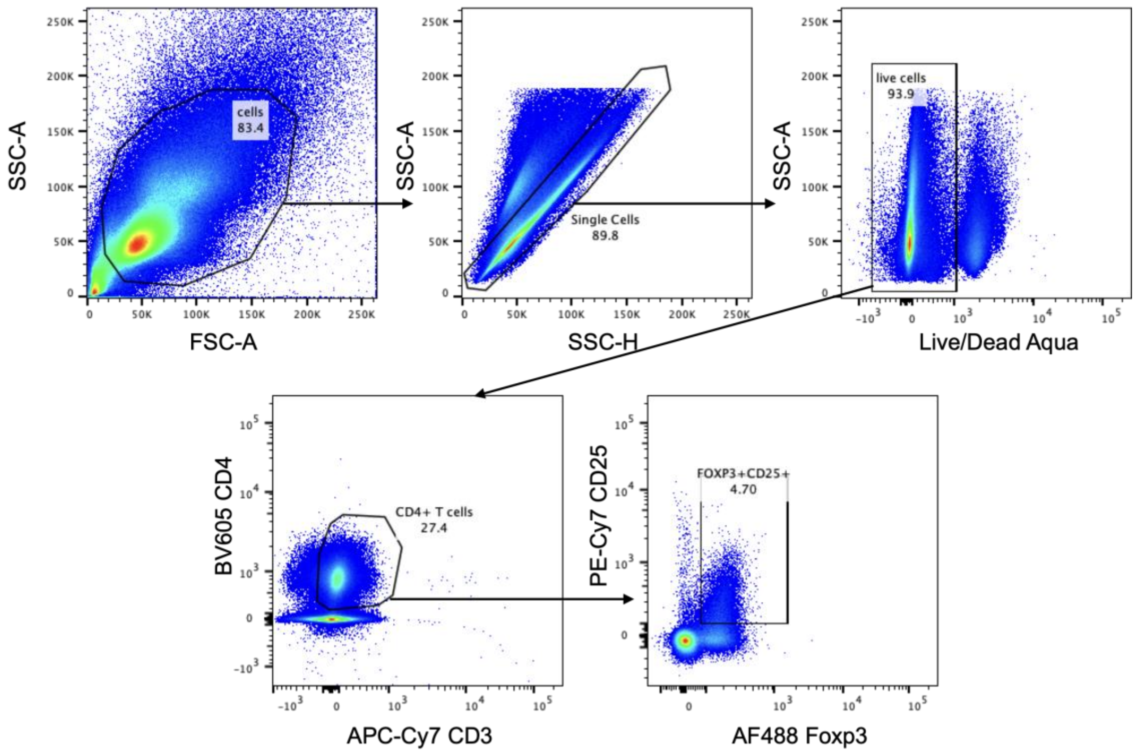


**Fig. S25.** Representative gating strategy for identifying $Foxp3^{+}CD25^{+}$ $CD4^{+}$ T cells present in spleen.

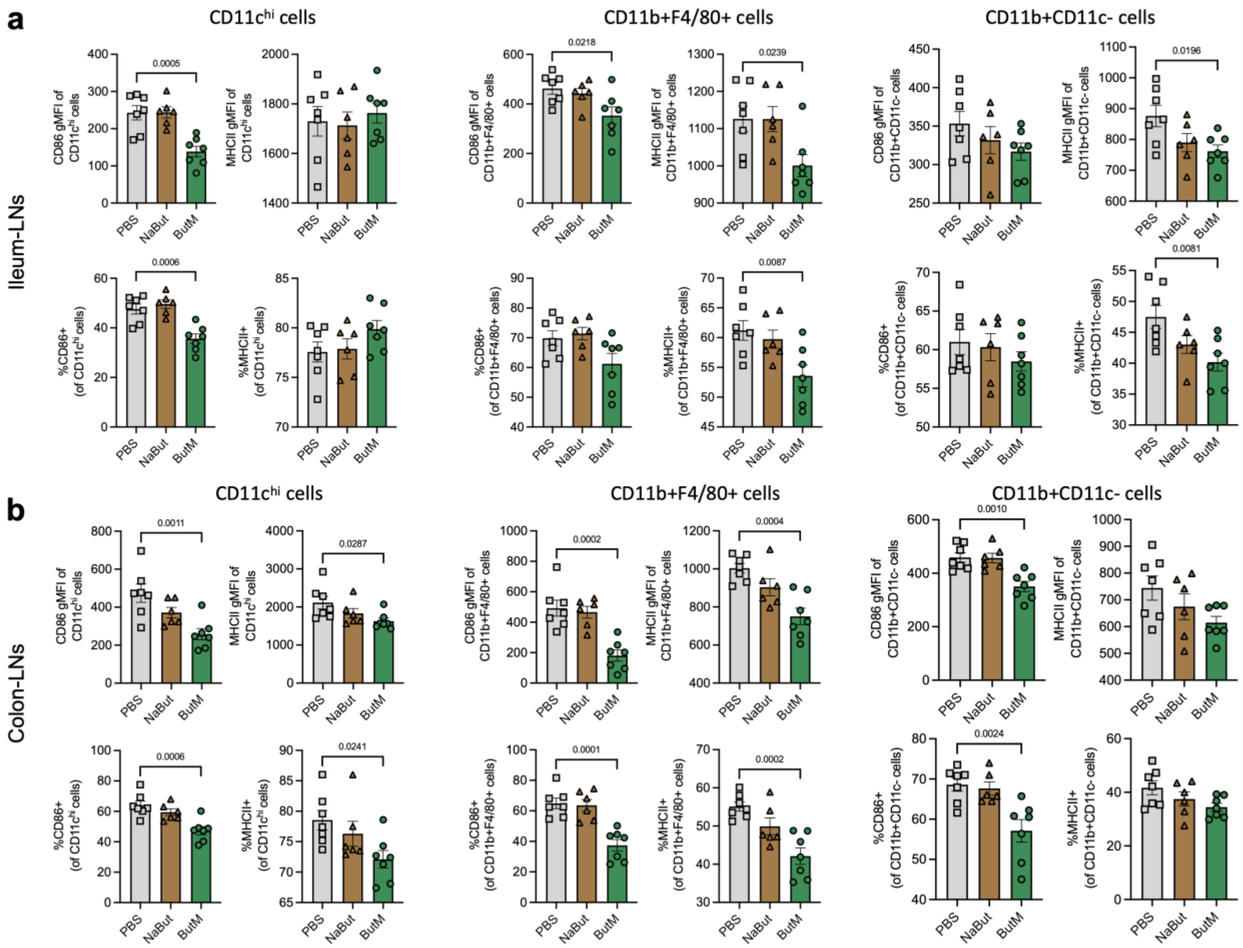


**Fig. S26.** Butyrate micelle treatment downregulated MHCII and the costimulatory molecule CD86 on the $CD11c^{hi}$, $CD11b^{+}F4/80^{+}$ and $CD11b^{+}CD11c^{-}$ cells in the ileum (**a**) and colon (**b**) draining LNs. The experimental schema is shown in **Fig. S24a**. Gating strategy is shown in **Fig. S27**. Data represent mean ± s.e.m. Data analyzed using one-way ANOVA with Dunnett's post-test.

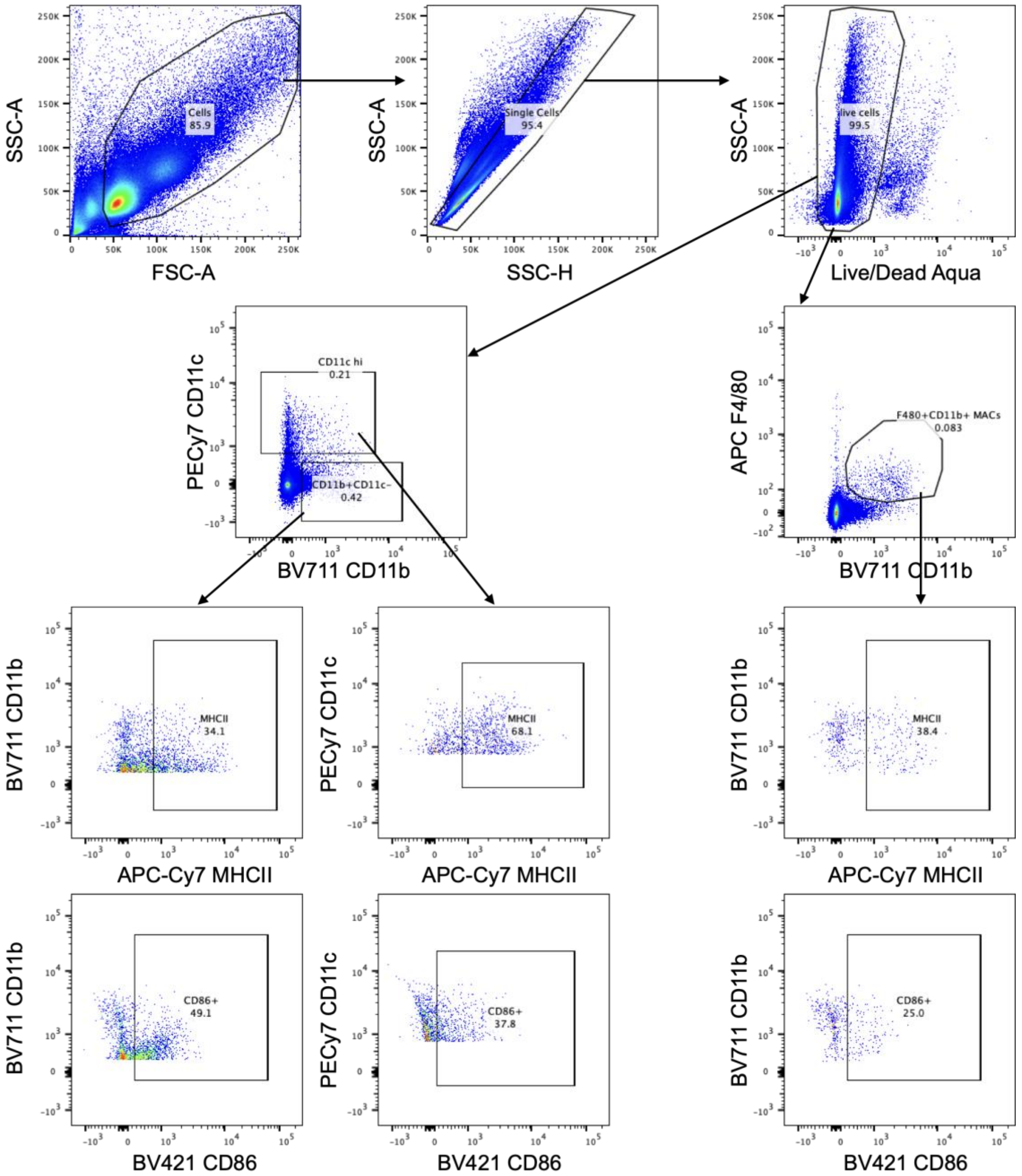


**Fig. S27.** Representative gating strategy for identifying CD11c$^{hi}$, CD11b$^{+}$F4/80$^{+}$, and CD11b$^{+}$CD11c$^{-}$ in the mesenteric LNs, and the MHCII$^{+}$ or CD86$^{+}$ cells in those cell subsets.

**Table S1.** Primer sequences for qPCR.

| Target | Primer sequence (5'->3') | Reference |
|---|---|---|
| *Clostridium Cluster XIVa* | Forward: AAATGACGGTACCTGACTAA | 73 |
| | Reverse: CTTTGAGTTTCATTCTTGCGAA | |

| **Primers to quantify total bacterial load** | **Primer sequence (5'->3')** | **Reference** |
|---|---|---|
| | 8F: AGAGTTTGATCCTGGCTCAG | 71 |
| | 338R: TGCTGCCTCCCGTAGGAGT | 72 |

| Target | Primer sequence (5'->3') | Reference |
|---|---|---|
| *Actb* | F: GGCTGTATTCCCCTCCATCG | 65 |
| | R: CCAGTTGGTAACAATGCCATGT | |
| *Defa3* | F: TCCTGCTCACCAATCCTCCAGGT | 42 |
| | R: CATATTGCGAACAATTTATTG | |
| *Defa5* | F: TCCTGCTCAACAATTCTCCAG | 42 |
| | R: CATATTGCAAACAATTTATTG | |
| *Defa20* | F: TCCTGCTCAACAATTCTCCAG | 42 |
| | R: CATATTGCAAACAATTTATTG | |
| *Defa21* | F: TCCTGCTCACCAATCCTCCAGGT | 42 |
| | R: CATATTGCAAACAATTTATTG | |
| *Defa22* | F: TCCTGCTCACCAATCCTCCAGGT | 42 |
| | R: CATATTGCAAACAATTTATTG | |
| *Defa23* | F: TCCTGCTCACCAATCCTCCAGGT | 42 |
| | R: CATATTGCGAACAATTTATTG | |
| *Defa24* | F: TGCTACTCACCAATCCTCCAGGT | 42 |
| | R: CATATTGCAAGCAATTTATTG | |
| *Defa26* | F: TCCTGCTCCCCAATCCCCCAGGT | 42 |
| | R: CATATTGCGGACAATTTATTG | |
| *Lyz1* | F: 5′-AAAACCCCAGGAGCAGTTAAT-3′ | 62 |
| | R: 5′-CAACCCTCTTTGCACAAGCT-3′ | |
| *Itln1* | F: '5'- ACCGCACCTTCACTGGCTTC-3' | 65 |
| | R: 5'- CCAACACTTTCCTTCTCCGTATTTC-3' | |
| *Slc10a2* | F: TTGCCTCTTCGTCTACACC | 42 |
| | R: CCAAAGGAAACAGGAATAACAAG | |
